# Structure of the mini-RNA-guided endonuclease CRISPR-CasΦ3

**DOI:** 10.1101/2021.06.27.449965

**Authors:** Arturo Carabias, Anders Fuglsang, Piero Temperini, Tillmann Pape, Nicholas Sofos, Stefano Stella, Simon Erlendsson, Guillermo Montoya

**Author notes:** Twelve Bio ApS, Ole Maaløes Vej 3, Copenhagen, 2200, Denmark (Stefano Stella).

## Abstract

CRISPR-CasΦ is a novel family of miniaturized RNA-guided endonucleases from phages ^1,2^. These novel ribonucleoproteins (RNPs) provide a compact scaffold gathering all key activities of a genome editing tool^2^. Here, we provide the first structural insight into CasΦ singular DNA targeting and cleavage mechanism by determining the cryoEM structure of CasΦ3 with the triple strand R-loop generated after DNA cleavage. The structure reveals the unique machinery for target unwinding to form the crRNA-DNA hybrid and cleaving the target DNA. The protospacer adjacent motif (PAM) is recognised by the target strand (T-strand) and non-target strand (NT-strand) PAM interacting domains (TPID and NPID). Unwinding occurs after insertion of the conserved α1 helix disrupting the dsDNA, thus facilitating the crRNA-DNA hybrid formation. The NT-strand is funnelled towards the RuvC catalytic site, while a long helix of TPID separates the displaced NT-strand and the crRNA-DNA hybrid avoiding DNA re-annealing. The crRNA-DNA hybrid is directed to the stop (STP) domain that splits the hybrid guiding the T-strand towards the RuvC active site. The conserved RuvC insertion of the CasΦ family is extended along the hybrid, interacting with the phosphate backbone of the crRNA. A cluster of hydrophobic residues anchors the RuvC insertion in a cavity of the STP domain. The assembly of the hybrid promotes the shortening of the RuvC insertion, thus pulling the STP towards the RuvC active site to activate catalysis. These findings illustrate why CasΦ unleashes unspecific cleavage activity, degrading ssDNA molecules after activation. Site-directed mutagenesis in key residues support CasΦ3 target DNA and non-specific ssDNA cutting mechanism. Our analysis provides new avenues to redesign the compact CRISPR-CasΦ nucleases for genome editing.

---

Competition between microbes and their invaders has driven the evolution of a wide catalogue of defence systems to prevent the attack of mobile genetic elements (MGEs). Among them, CRISPR constitutes a type of adaptive immunity achieved by CRISPR-associated nucleases (Cas) and CRISPR RNAs (crRNAs) that assemble effector ribonucleoprotein complexes, which are guided by the crRNA to recognise and cleave complementary DNA (or RNA) for interference ^3–5^. CRISPR-Cas nucleases have been extensively used as tools for genome editing ^6,7^. The redesign of their guide RNA to target specific DNA sites, as well as the manipulation of the protein scaffold ^8,9^, has provided a powerful method for genome modification in biomedical and biotechnological applications^10,11^.

Although ubiquitously diversified among prokaryotes, CRISPR systems were also identified in the genome of bacteriophages ^1^. Recently, a new Class 2 family of CRISPR nucleases named CasΦ proteins, also known as Cas12j, were found in the biggiephage clade of “huge” phages^2^. CasΦ proteins share a sequence identity lower than 7% with other CRISPR nucleases^2^ and display sequence and structural homology only in their RuvC domain with Class2 type V members^2^. CasΦ RNPs generate a staggered DNA double strand break (DSB) and unleash unspecific ssDNA cleavage after activation with a ssDNA molecule complementary to the crRNA, as other members of the Class 2 type V nucleases ^12^. In addition, the RuvC catalytic site of CasΦ1 and 2 also processes the precursor crRNA (pre-crRNA)^2^. CasΦ endonucleases recognise protospacers with a minimal T-rich PAM, and their small size (700-800 residues) together with the lack of a trans activation crRNA (trac-crRNA) to build the functional RNP, make CasΦ a unique family of miniaturized RNA-guided nucleases (Fig. 1a-b, Extended Data Fig.1).

**Figure 1.**
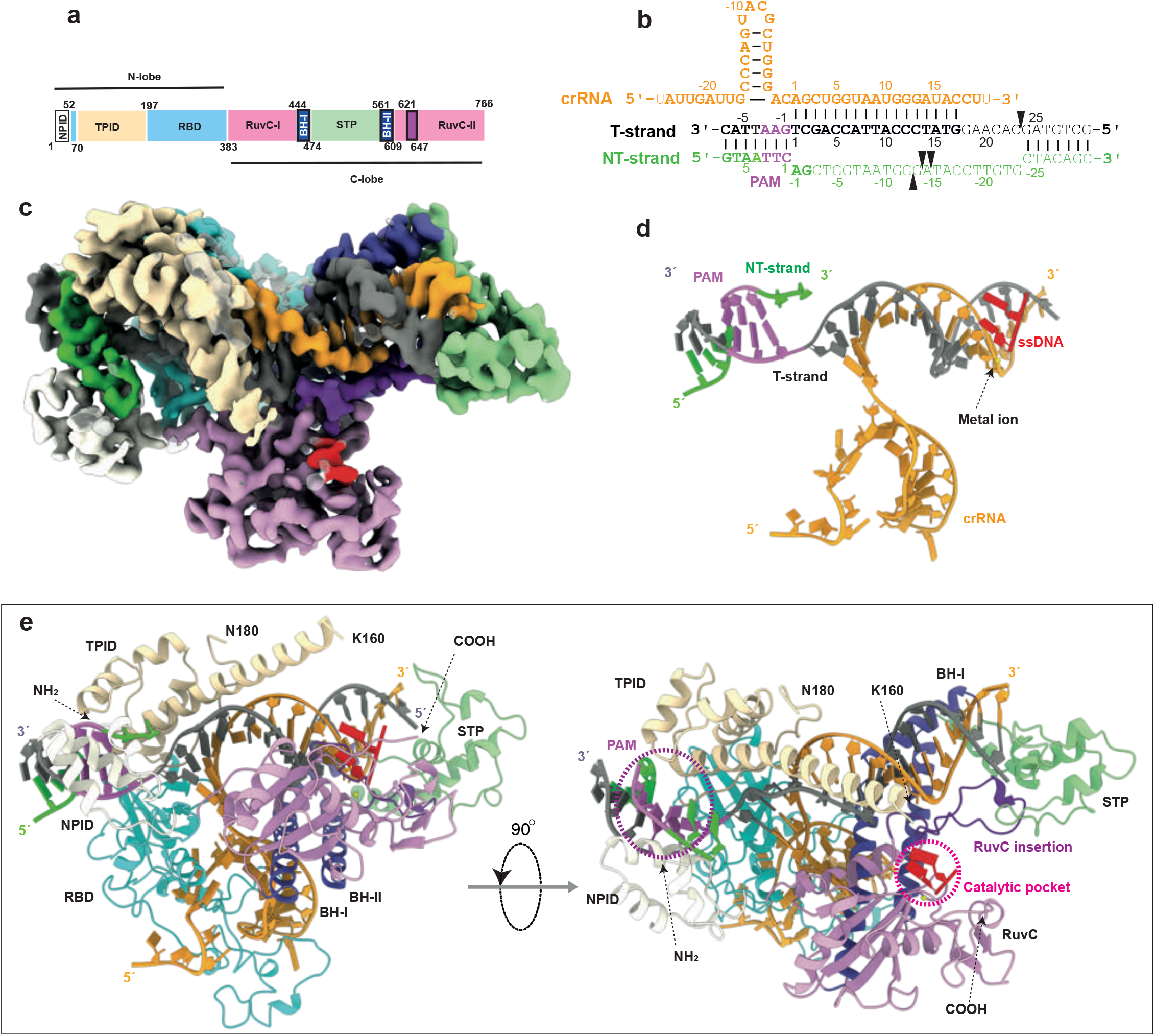
Cryo-EM structure of CasΦ3 endonuclease R-loop complex after target DNA cleavage. **a)** Domain architecture of CasΦ3 comprising the T-strand and NT-strand PAM interacting domains (TPID, NPID), the RNA-handle binding domain (RBD), the bridge helices (BH-I and BH-II), the RuvC domain including the insertion (amino acids 621-647) and the stop (STP) domain. **b)** Schematic diagram of the R-loop formed by the crRNA and the target DNA. Triangles represent phosphodiester cleavage positions in the T- and NT-strands; the light font nucleotides represent those not visualized in the structure. **c)** cryo-EM map of the CasΦ3/R-loop complex at 2.7 Å resolution. The map is coloured according to each domain as in Fig. 1a. (Extended Data Fig. 3-5, Extended Data Table I). **d)** View of the R-loop structure and 2 nucleotides and the divalent metal ion in the catalytic site (polypeptide omitted for clarity). **e)** Overview of the CasΦ3–RNA–target-DNA ternary complex.

CRISPR-Cas effector complexes are harnessed *in vitro* and *in vivo* for genome editing approaches, but specially the latter is limited by delivery problems, which is one of the main unmet needs in the field ^11^. Adeno-associated viral vectors (AAV) are commonly used for gene delivery. Yet, packaging of the genes coding for CRISPR-Cas effector complexes into an AAV vector is challenging due to its limited capacity, thus leaving little space for the insertion of additional regulatory elements. Recently, CasΦ enzymes have been shown to mediate genome editing in mammalian and plant cells^2^ expanding our *repertoire* of genome manipulation tools. The small size CasΦ RNPs can improve our genome editing approaches by alleviating the packing problems in the AAV vectors used for delivery ^11^. However, questions regarding the detailed molecular mechanism of target DNA recognition, unzipping and subsequent cleavage by CasΦ nucleases remain unanswered, as no structural information is available.

## CasΦ3/R-loop structure determination

We reconstituted and characterized a functional CasΦ3-crRNA complex (Extended Data Fig. 2, Methods) and determined the structure of the enzyme after severing a target dsDNA by cryo-EM (Fig. 1, Extended Data Fig. 3-5, Methods). Heterogeneous refinement resulted in several conformations of the complex. The predominant class yielded a map at a resolution of 2.7 Å, which was used to build the model of the CasΦ3/R-loop structure. The high flexibility observed in the second predominant class precluded building a complete model but revealed the flexible regions and the conformational heterogeneity of the complex (Extended Data Fig. 3-5, Extended Data Table I, Supplementary Video 1). The CasΦ3/R-loop structure represents a snapshot of the endonuclease-product complex after substrate cleavage (Fig. 1c-e), revealing the critical residues for PAM recognition, target DNA unwinding and cleavage, and thereby providing detailed atomic information for the redesign of this novel family of genome editing tools.

## CasΦ3 biochemical characterisation

CasΦ3 generates an overhang of 9-11 nucleotides by cleaving a specific target DNA at different phosphodiester bonds (Fig. 1b, Extended Data Fig. 2a). A collateral effect of its specific cleavage is the release of indiscriminate ssDNA degradation^2^, which is triggered by the T-strand provided as target dsDNA or as a ssDNA activator complementary to the crRNA (Extended Data Fig. 2b-c). In both cases, indiscriminate CasΦ3 cleavage is unleashed when a minimal 12-to 13-nt crRNA-DNA duplex is assembled. The structure suggest that the differences observed with activators longer than 18-nt can be attributed to the presence of the R-loop disturbing the entrance of the unspecific ssDNA substrate in the catalytic site (Fig. 1d-e, Extended Data Fig. 2d, see below). The activity of the endonuclease was tested in the presence of Mg^2+^ and other divalent metal ions (Extended Data Fig. 2e). The assay revealed that CasΦ3 supports catalysis in the presence of Mn^2+^, Fe^2+^, Co^2+^, and Ni^2+^ resulting in different cleavage patterns. CasΦ3 cleavage activity was saturated when the endonuclease/target-DNA ratio was nearly equimolar, suggesting the slow dissociation of the enzyme from the PAM-proximal cleavage product, as observed in other RNA-guided nucleases ^13,14^ (Extended Data Fig. 2f). In addition, removing the last 39 residues of the C-terminus, which were not visualized in the structure, decreased CasΦ3 activity. However, the enzyme conserved a substantial catalytic activity, suggesting that CasΦ family members can be further miniaturized (Extended Data Fig. 2g-h).

## Overall structure of the CasΦ3/R-loop complex

The CasΦ3/R-loop complex does not present the classical bilobal architecture observed in other type V effector complexes. The R-loop displays a T shape with the crRNA/DNA hybrid and the crRNA handle forming the horizontal and vertical bars, and the protein domains wrapping around the nucleic acids (Fig. 1d-e). The handle of the crRNA is stabilized by the strictly conserved R338 which interacts with C-1 and U-18 and the neighbouring non-Watson-Crick base pair interaction between G-17 and A-2 (Extended Data Fig. 6). The PAM-distal and PAM-proximal regions of the heteroduplex are recognized by the N- and C-terminal regions of the polypeptide (Fig. 1d-e), which are connected by a 15-residue loop (380-395). Each region comprises around half of the size of the protein and they are separated by the long handle of the crRNA on the T-shape assembly. The N-terminal region comprises the T-strand and NT-strand PAM interacting domains (TPID, NPID) and the RNA-handle binding domain (RBD), while the C-terminal consists of the catalytic RuvC and the stop (STP) domains (Fig. 1a). The RuvC domain is split into RuvC-I and RuvC-II by the insertion of the STP domain, which is connected to the catalytic domain by two long bridge helices, BH-I and BH-II. Additionally, the RuvC-II subdomain presents a characteristic insertion, which is conserved in all the known members of the CasΦ family except CasΦ7 (Fig. 1, Extended Data Fig. 1). This N- and C-terminal physical separation is also functional, as the RNP assembly, PAM recognition and unwinding reside in the N-terminal region, while the crRNA/T-strand hybrid assembly and catalysis of the target DNA are performed by the C-terminal section of the polypeptide. Therefore, the PAM binding site is ~55Å away from the RuvC nuclease active site.

The target DNA cleavage yields a triple strand R-loop with the T-strand hybridized to the crRNA (Fig. 1b, d), while the dissociated PAM NT-strand is directed towards the RuvC catalytic pocket (Fig. 2a). The NT-strand nucleotides −1 to −2 upstream of the PAM were built in the density but the high flexibility on the distal end of the NT-strand precluded visualization of the rest of the nucleotides, as shown for Cas9^15^ and Cas12a^13^. Nevertheless, the backbone of the NT-strand is observed at low contour level in the cryo-EM maps, suggesting the path followed by the DNA to the RuvC catalytic pocket (Figure 2a, Extended Data Fig. 7). Interestingly, two nucleotides, modelled as purines, were observed in the RuvC pocket in complex with Ni^2+^ as a by-product of the phosphodiester hydrolysis (Fig. 1c-e, 2a, Methods). To determine to which strand these nucleotides belong, we performed a binding assay after cleavage with different labelled target DNA, revealing that these nucleotides originate from the NT-strand (Extended Data Fig. 7c).

**Figure 2.**
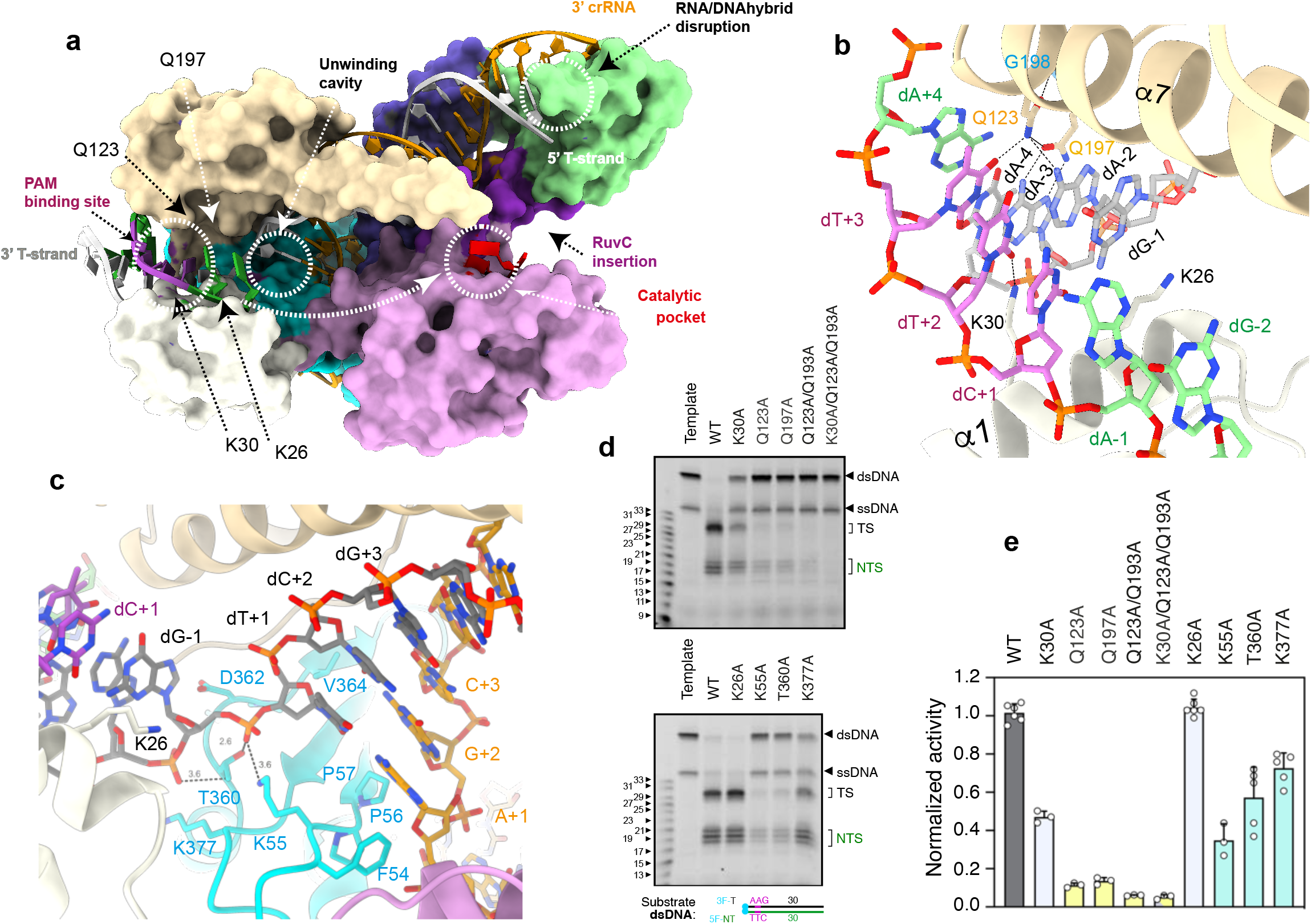
CasΦ3 PAM recognition, uncoupling of the Watson–Crick dA-1:dT+1 pair and unzipping. **a**) Surface representation of CasΦ3–R-loop complex. The white dashed arrow shows the predicted path of the NT-strand to the DNA nuclease site after dG-2 (Extended Data Fig. 6). **b)** Detailed view of the PAM nucleotides recognition and the dsDNA unwinding depicting the interactions of the conserved K26, K30, Q123 and Q197 residues (Extended Data Fig. 5c). **c**) Zoom of the dT+1/dA-1 pair uncoupling, phosphate inversion and unzipping (Extended Data Fig. 5d). Black dashed lines in b) and e) represent polar interactions between 2.2 and 3.2 Å. **d)** Representative dsDNA cleavage assays using CasΦ3 wild type (WT) and mutants. Oligonucleotides 3F-T-AAG-30 and 5F-NT-TTC-30 were used as substrate (Extended Data Table II). T-strand (TS) and NT-strand (NTS) products are marked. Each experiment was repeated three to six times. **e)** Quantification of the activity based on the cleavage experiments as shown in d). Bars represent mean ± s.d.

## PAM recognition

PAM recognition is an important aspect of DNA targeting by CRISPR-Cas nucleases, as it is a prerequisite for target DNA identification, strand separation and crRNA–target-DNA heteroduplex formation^16^ before cleavage. CasΦ3 is reported to recognize a 5′-TTN-3′ PAM sequence in the NT-strand ^2^. Our structure shows that PAM recognition in CasΦ3 is achieved by a combination of interactions in both strands by the TPID and NPID domains (Fig. 2b, Extended Data Fig. 5c, 6). The positively charged side of helix α1 (S21 to A34, Extended Data Fig. 1) in the NPID is inserted in the minor groove at an angle of 45° with respect to the dsDNA longitudinal axis, thus facilitating the unwinding of the dsDNA. Two conserved lysines, K26 and K30, interact with the NT-strand. K30 makes specific contacts with dT+2, while K26 is placed inside the dsDNA to disrupt Watson-Crick base coupling, displacing the NT-strand and promoting separation (Fig. 2b-c, Extended Data Fig. 5c-d). On the other side of the PAM recognition cleft, Q123 in the TPID builds an intricate network of polar interaction with dA-3, dA-2 in the T- and the dT+3 in the NT-strand (Fig. 2b, Extended Data Fig. 6). The neighbouring G198 amide contacts the carbonyl of Q123, anchoring the side chain in a conformation favouring the contacts with these bases. In addition, the side chain of Q197 interacts with Q123 and hydrogen bonds with dA-3.

The Q123A and Q197A mutations present ~90% activity reduction, while the K30A mutant reduces cleavage ~55%. The triple mutant activity is similar to the Q123A/Q197A mutant, indicating the pivotal role of the glutamines in PAM recognition, as the addition of the K30A mutation does not display a further reduction (Fig. 2d-e). The K26A mutant activity is not affected, suggesting that the insertion of the α1 helix is sufficient to unzip the dsDNA. All the mutants involved in PAM recognition do not change the cleavage pattern of the dsDNA target (Fig. 2d-e). Both the wild type and the mutants did not cleave target dsDNAs with different PAM sequences or in the absence of PAM, underscoring the selectivity of the PAM interaction network formed by Q123A, Q197A and K30A (Extended Data Fig. 8a). In addition, we observed that the unspecific ssDNA catalysis is also fully activated in the presence of dsDNA containing the PAM, thus, suggesting that after PAM recognition crRNA/DNA hybrid assembly activates catalysis (Extended data Fig. 8b). Finally, to assess the role of the PAM complementary bases in the T-strand, we triggered the unspecific activity of CasΦ3 using ssDNAs activators mimicking the T-strand with different PAM sequences. The assay showed that the PAM complementary 3’-AAG-5’ sequence and an activator without PAM, fully released phosphodiester hydrolysis, while other PAMs promoted activation to different levels. This experiment suggests that the assembly of the proper hybrid unleashes the catalytic activity, while activators containing regions that partially hybridize with the crRNA display lower cleavage (Extended Data Fig. 8b).

Collectively, our analysis suggests that the well-conserved Q123 and Q197 residues, which interact with the PAM in the major groove of the target DNA, play an essential role in recognition. The direct base readout in the PAM region of CasΦ nucleases combine interactions of the TPID and NPID with both strands of the target DNA. However, the interactions of the TPID with the T-strand seem to have an important role in PAM discrimination. This is a singular property of the CasΦ family, as other CRISPR-Cas nucleases perform PAM scanning by interacting preferentially with the NT-strand ^17,18^.

## Target DNA unwinding

Overlaying with the first uncoupled base pair upstream the PAM, the TPID, NPID and the antiparallel β-sheet composed of the β1, β6 and β7 strands of the RBD domain (Extended Data Fig. 1), build a cavity where unwinding and the initial crRNA/T-strand hybridisation occurs (Fig. 2c). This cavity is flanked on the C-terminal region by the BH-I helix and the RuvC domain. The well-conserved F54, K55, P56, P57, P363, T360, G361, D362 and V364 organize the cavity combining acidic and hydrophobic residues facilitating the Watson-Crick base pairing of dT+1 and A+1 in the T-strand and the seed of the crRNA (Fig. 2c, Extended Data Fig. 5d, 6). In addition, the backbone phosphate group of dG-1 is recognized by the side chain of the T360, K55 and the main chain of Y376. This interaction results in the rotation of the phosphate group (Fig. 2c, Extended Data Fig. 6), facilitating base pairing between dT+1 and A+1 in the crRNA as observed in Cas9^19^ and Cas12a complexes^13,20–23^. The neighbouring K377A mutation led to ~20% decrease in the activity, but the T360A and the K55A mutations displayed a reduction of 50% and 60%, highlighting the importance of these residues for phosphate inversion and hybrid formation (Fig. 2d-e). The long helix α7 in the TPID directs the crRNA/T-strand hybrid into the “nest” formed by the BH-I and II helices and the RuvC insertion, and detaches the hybrid from the NT-strand preventing a possible reannealing of the target DNA. The area where the hybrid rests is flanked by the catalytic RuvC and STP domains, which disrupts the crRNA/T-strand hybrid as a vessel bulb bow (Fig. 3a). An antiparallel β-sheet formed by β11 and β12 splits the Watson-Crick base coupling after the dG-17:C+17 pair; thus, limiting the hybrid length to 17 nucleotides in agreement with cleavage experiments testing the efficiency of the spacer length ^2^. The aromatic ring of F538 in β11 initiates the hybrid unzipping (Fig. 3a). The 3’-phosphate of the crRNA is guided to the back side of the domain, where C+17 and U+18 are accommodated by a combination of basic (R535, R547) and hydrophobic residues (M500, L555), and the 5’-phosphate of the T-strand is directed to the other side of the protein where the RuvC catalytic pocket is located.

**Figure 3.**
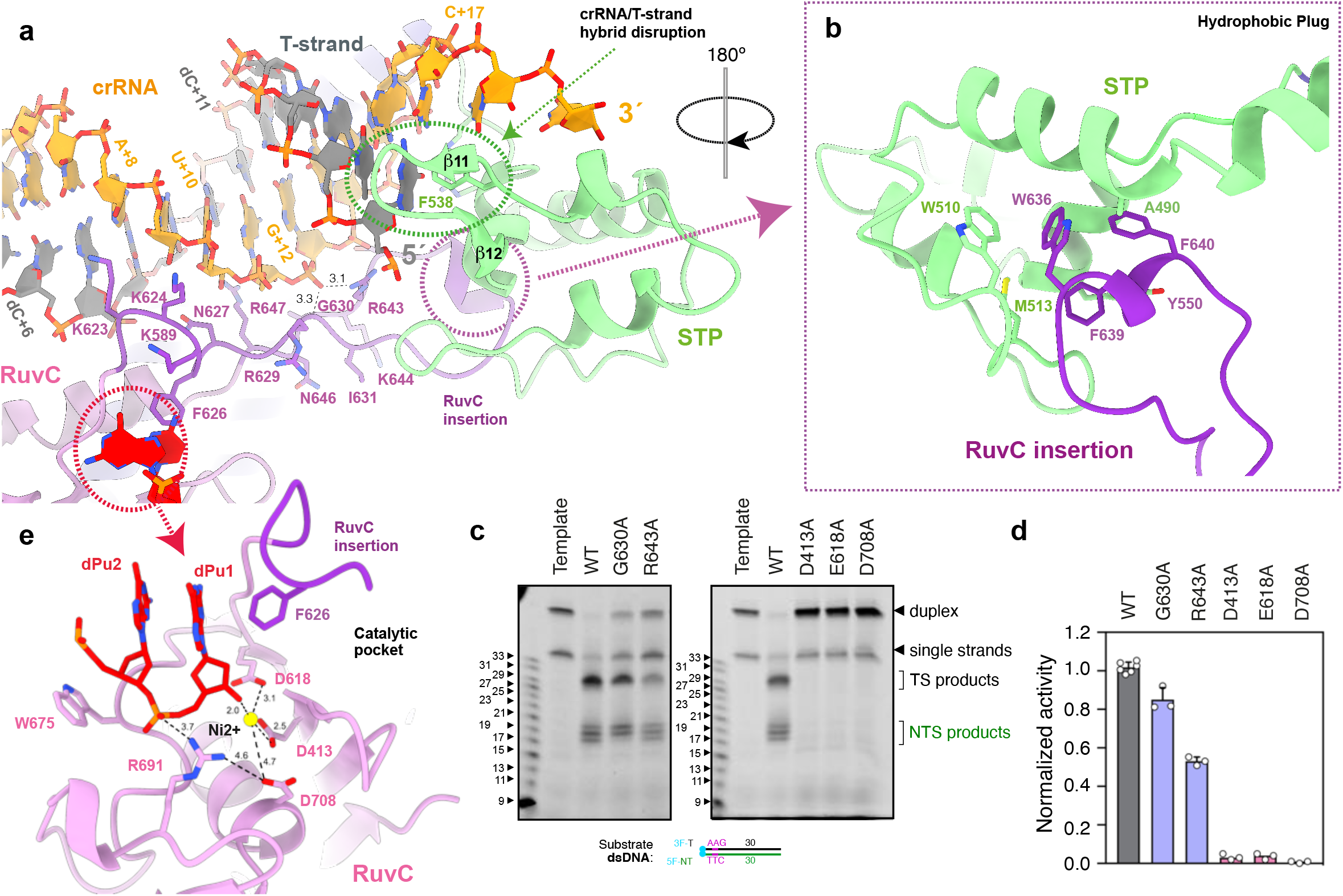
Assembly of the crRNA/DNA hybrid activates catalysis in the RuvC pocket. **a)** View of the hybrid showing the interaction of the crRNA with residues in the RuvC insertion (Extended Data Fig. 5e). **b)** Inset depicting the hydrophobic interaction between the “plug” of the RuvC insertion and the and cavity of the STP domain (Extended Data Fig. 5f). **c)** Representative dsDNA cleavage assays using CasΦ3 wild type (WT) and mutants. Oligonucleotides 3F-T-AAG-30 and 5F-NT-TTC-30 were used as substrate (Extended Data Table II). T-strand (TS) and NT-strand (NTS) products are marked). Each experiment was repeated three times. **d)** Quantification of the activity based on the cleavage experiments as shown in c). Bars represent the mean ± s.d. **e)** Detailed view of the RuvC catalytic site containing a dinucleotide and a divalent metal. The D708 side chain and the associated distances are shown for visualization purposes and (Extended Data Fig. 5g). Black dashed lines in a) and e) represent polar interactions between 2.0 and 3.5 Å.

## Catalytic activation

The RuvC insertion runs alongside the crRNA strand of the hybrid, making multiple contacts with its phosphate backbone from U+9 to G+13 (Extended Data Fig. 6, 5e), and the turn at the tip of the insertion is anchored in the back side of the STP domain by hydrophobic interactions (Fig. 3b, Extended Data Fig. 5f). This arrangement and the activity assays (Extended Data Fig. 2b-c, 8c-d), suggest that the assembly of the crRNA/DNA hybrid could trigger conformational changes in the RuvC insertion that activate catalysis by making the active pocket available for the ssDNA substrate. The monitoring of the unspecific cleavage of ssDNA substrate using activators of different length (Experimental Data Fig. 2b-c), shows that the unspecific activity of CasΦ3 is fully released when the activator’s length allows the formation of a 12-nt crRNA/DNA hybrid or longer, supporting the notion that a certain hybrid length is needed to activate catalysis. The conserved G630 and R643 are key residues, as they arrange a network of polar interactions with the phosphate of G+12, resulting in a special arrangement of the connections joining the hydrophobic “plug” composed by the conserved W636, F639 and F640 residues in the tip of the insertion (Fig. 3a-b). We hypothesize that the assembly of the hybrid would promote the observed conformation of the RuvC insertion, which is anchored by the plug in the cleft of the STP domain composed by A490, W510, M513 (Fig. 3b). The stabilisation of this conformation by the hybrid would pull the STP domain towards the catalytic site, placing the T-strand in the active site with the proper 5’−3’polarity. Mutations in the hydrophobic plug and STP cleft residues rendered CasΦ3 insoluble, highlighting the importance of this conserved interaction in the CasΦ family.

To analyse our hypothesis, we generated the R643A and R630A mutants. The activity of the R643A mutant is reduced to ~50%, while the G630A exhibited a minor decrease of ~20% (Fig. 3c-d), which could be explained by the fact that G630 contributes to the polar network with the G+12 phosphate through its main chain. Interestingly the phosphodiester cleavage pattern of G630A displays only two sites in the NT-strand. Next, we tested our assembly activation hypothesis examining the unspecific ssDNA degradation of the PAM, unwinding and the R643A mutants with an 18-nt ssDNA activator without PAM (Experimental Data Fig. 8c-d). All the PAM and unwinding mutants display full enzymatic activity while the R643A presents a 50% reduction. When the same assay was performed using a dsDNA containing the PAM for activation, none of the PAM mutants displayed noticeable activity, while the R643A still presented 50% reduction. These results support the proposed model, as the ssDNA activator without PAM should skip recognition and unwinding, thus directly hybridising with the crRNA and triggering full enzymatic activity in all the PAM and unwinding mutants. However, when the PAM is present in the target dsDNA these variants displayed reduced cleavage as PAM recognition and unwinding are compromised, in agreement with the results observed in specific dsDNA cleavage assays (Fig. 2 d-e). By contrast the activity of the R643 mutant, which is essential in monitoring crRNA/DNA assembly, is affected independently of the presence or absence of PAM in the DNA used for activation (Extended Data Fig. 8c-d). Therefore, PAM recognition, DNA unwinding and RuvC catalytic activation are linked in the presence of a target dsDNA, while the catalytic activation of RuvC can omit PAM recognition if a suitable ssDNA is provided to the enzyme. Furthermore, mutations in the RuvC insertion do not only affect the enzyme activity, but they could potentially change the cleavage pattern as observed in the case of the G630A mutant.

## DNA cleavage

The RuvC domain of CasΦ nucleases belong to the retroviral integrase superfamily that displays a characteristic RNaseH fold. The two nucleotides from the NT-strand in the catalytic CasΦ3 pocket are associated with the conserved E618 and D413 (Fig. 3e). The density did not allow base identification, and either dA or dG could be modelled. We built two guanines with a 5’−3’ polarity and a Ni^2+^ ion (Methods), in agreement with the number of nucleotides in the cleavage products and the purine rich sequence in that position (Fig.1b, 3e, Extended Data Fig. 6, 7). Therefore, the length of the DNA after DSB generation could permit that the cleaved NT-strand remains associated with the catalytic centre and may disturb the entrance of the T-strand delaying its catalysis, as previously observed ^2^ (Extended Data Fig. 2g). A second metal atom, modelled as Zn, is coordinated by 4 conserved cysteines, similarly to Cas12f ^24^ and Cas12g ^25^ (Extended Data Fig. 9a-b). This section of RuvC includes the conserved R691 3.7 Å away from the dinucleotide. This residue could facilitate the positioning of the phosphodiester backbone in the catalytic pocket (Fig. 3e). However, the rest of this region is different to the target nucleic acid-binding (TNB) domain in Cas12f and Cas12g (also known as the Nuc domain for Cas12a and Cas12b and the target-strand loading domain for Cas12e), as it displays a different structure that does not contain the helical regulatory lid motif.

RuvC domains introduce 5′-phosphorylated cuts and involve three acidic amino acids^26^ and two divalent metal ions ^27^. The E618 and D413 carboxylate amino acids are important catalytic residues, and the E618A and D413A mutations abolish CasΦ3 activity (Fig. 3c-e). Both residues are predicted to coordinate the metal ions that activate the nucleophile and stabilize the transition state and the leaving group. In our structure, E618 and D413 coordinate the metal and the backbone of the dinucleotide (Fig. 3e, Extended Data Fig. 6). The side chain of D708, which is predicted to act as the third catalytic residue, is not observed due to electron irradiation ^28^. This active-site residue has been shown less critical than the other carboxylates in other RuvC domains, and substitutions of this amino acid to Asn or His lead to only partial loss of cleavage ^29,30^. However, the D708A mutation abrogates activity (Fig. 3c-e). Structural comparisons using DALI with other RuvC domains, including CRISPR-Cas proteins, support a two metal ion mechanism (Extended Data Fig. 9). Interestingly, we cannot observe differences with the RuvCs of CasΦ1 and 2 that could explain why CasΦ3 is unable to cleave, and thereby process, its own crRNA, as the sequence homology in this domain is high within the CasΦ family (Experimental Data Fig. 1).

## Discussion

These findings suggest a working mechanism for CasΦ3 that explains the enzymatic activation when target dsDNA or ssDNA activator is used (Fig. 4). CasΦ3 will bind the target DNA, inducing the interaction with the TPID, thus allowing scanning of PAM. Binding to PAM fosters the stabilisation of NPID on the NT-strand domain and the insertion of the α1 helix. Then, the DNA duplex is unzipped and coupling between the T-strand and the crRNA occurs. The α7 helix separates the crRNA/T-strand hybrid away from the NT-strand, which is guided to the active site in RuvC with a 5’−3’ orientation. The STP domain splits the hybrid and orients the T-strand towards the RuvC active site. The length of the hybrid is monitored by the RuvC insertion whose conformational change induced by the crRNA/DNA hybrid pulls the STP domain towards the catalytic site activating phosphodiester hydrolysis on the NT-, and guiding the T-strand for catalysis to the active pocket for cleavage. Catalysis proceeds to generate the overhang on the target DNA. The unspecific ssDNA cleavage activity would be triggered by the same mechanism after crRNA/activator DNA assembly.

**Figure 4.**
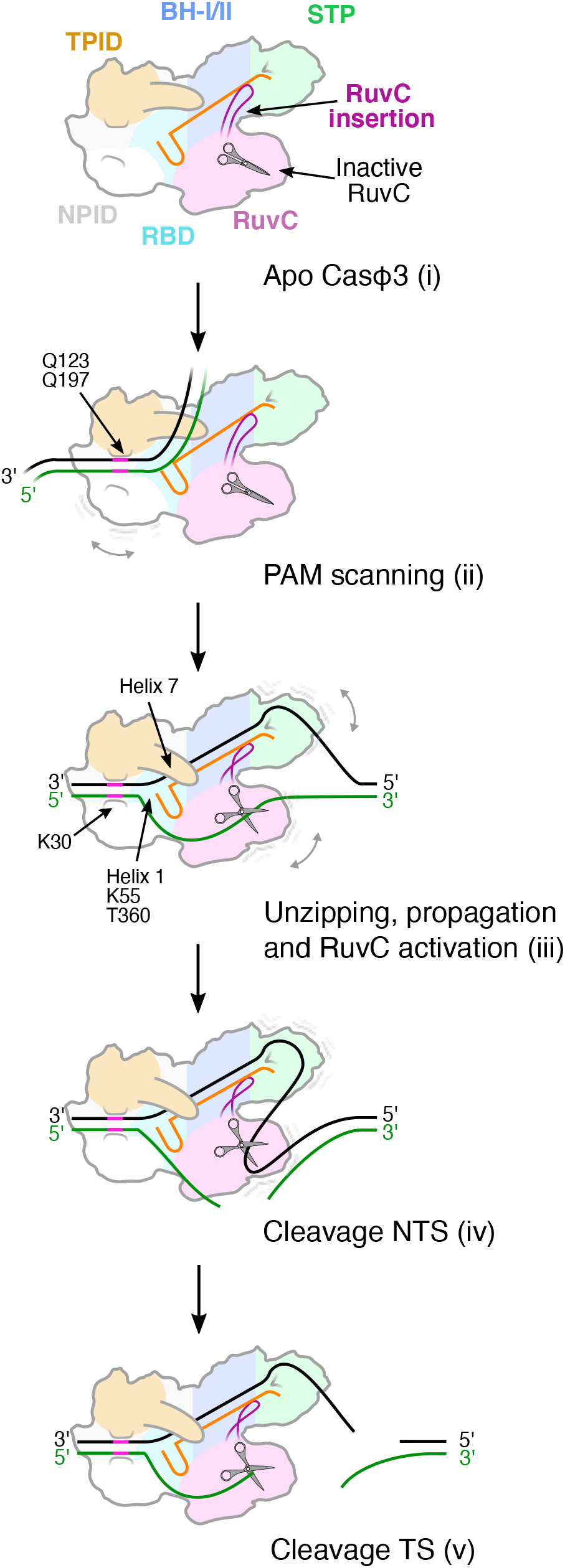
Model of CasΦ3 PAM-dependent DNA recognition, unwinding and cleavage. Cartoon model depicting the stages of CasΦ3 nuclease staggered target DNA cleavage.

This work provides a detailed molecular understanding into the unique CasΦ endonuclease family. CasΦ mediate genome editing in eukaryotic cells^2^ and conserve the properties of other CRISPR-Cas nucleases in a tiny RNP. Therefore, they could mitigate packing restrictions in the AAV vectors used for delivery ^11^. Our study paves the way for the rational redesign of CasΦ endonucleases widening the CRISPR-Cas toolbox.

## Acknowledgements

We thank the Danish Cryo-EM National Facility in CFIM at the University of Copenhagen for support during cryo-EM data collection supported by grant NNF0024386. Data processing has been performed at the Computerome, the Danish National Computer for Life Sciences. G.M. is a member of the Integrative Structural Biology Cluster (ISBUC) at the University of Copenhagen. SE is funded by the Novo Nordisk Foundation (Grant NNF17OC0030788). G.M. is part of the Novo Nordisk Foundation Center for Protein Research (CPR), which is supported financially by the Novo Nordisk Foundation (grant NNF14CC0001). This work was also supported by the cryoNET (grant NNF17SA0030214), and Distinguished Investigator (NNF18OC0055061) grants to G.M.

## Author Contributions

AC, AF, SS and GM designed the biochemical experiments. ACR, AF and PT set up the purification protocol. AC created the mutants and performed biochemistry experiments using CasΦ3 mutants. PT purified CasΦ3 WT and mutants. AF performed most of the biochemistry analysis. AC, AF and GM analysed the biochemical data. AC and AF prepared the cryo-EM sample, EM grids and collected the cryo-EM images. TP and NS helped with sample preparation, data collection and advised in the initial stages of processing. SE helped with the Warp pipeline, and AC performed the rest of the Cryo-EM processing and built the structure. AC, AF and GM proceeded with cryo-EM map and structure analysis. The global results were discussed and evaluated with all authors. GM coordinated and supervised the project and wrote the manuscript with input from all the authors.

## Author Information

Guillermo Montoya and Stefano Stella declare that they are co-founders of Twelve Bio. A patent application has been filed relating to this work.

## Methods

### Plasmid preparation, protein expression and purification

CasΦ3 cDNA was synthetized and cloned with a C-terminal hexahistidine (His)-tag into pET-21 vector (Genewiz) (Extended Data Table II). CasΦ3 mutants were generated with the In-Fusion cloning kit (Takara) using the primers specified in Extended Data Table II. To generate CasΦ3-ΔCT, a TEV cleavage site (ENLYFQG) was generated after the residue M726. His-tagged CasΦ3 was expressed from pET-21 in *E. coli* BL21 pRARE cells. *E. coli* cultures were grown at 37° C in liquid Terrific Broth (TB) medium with 34 mg/l chloramphenicol and 100 mg/l ampicillin to an optical density at 600 nm of ~ 0.8. Overexpression of proteins was induced with 150 nM of IPTG for 16h at 16°C. Cells were harvested by centrifugation and resuspended in lysis buffer (50 mM HEPES pH7.5, 2M NaCl, 5 mM MgCl_2_, 1 tablet of Complete Inhibitor cocktail EDTA Free (Roche) per 50◻ml, 50◻U/ml Benzonase, 1◻mg/ml lysozyme). Lysis was completed by one freeze-thaw cycle and sonication. Cell extract was diluted to a final salt concentration of 500 mM, and high-speed centrifuged (10,000 x g, 45 min) to separate the soluble fraction from the insoluble fraction and the cell debris. The soluble fraction was loaded into a 5 ml HisTrap FF Crude column (Cytiva) equilibrated in buffer IMAC-A (20 mM HEPES pH7.5, 500 mM NaCl, 20 mM Imidazole), and bound proteins were eluted by stepwise increase of the imidazole concentration with buffer IMAC-B (20 mM HEPES pH7.5, 200 mM KCl, 500 mM Imidazole). CasΦ3 proteins eluted at ~150 mM Imidazole. In the case of CasΦ3-ΔCT, the C-terminal segment (residues 727-766) was cleaved by incubating the protein with 0.3 mg TEV protease in TEV buffer (20 mM HEPES pH 7.5, 150 mM NaCl, 1 mM EDTA, 0.5 mM TCEP) for 16 h at 4 °C. Fractions containing CasΦ3 were pooled, concentrated and further purified by size exclusion chromatography (SEC) using a HiLoad 16/600 Superdex 200 column (Cytiva) equilibrated in SEC buffer (20 mM HEPES pH7.5, 500 mM KCl, 0.5 mM TCEP). Fractions containing pure protein were pooled, concentrated to 5-10 g/L, flash-frozen in liquid nitrogen and stored at −80 °C.

### Cleavage assays

Fluorescein (FAM)-labeled DNA oligonucleotide at 5’ or 3’ ends, unlabeled DNA and RNA oligonucleotides were purchased from Integrated DNA technologies (IDT) (Extended Data Table II). dsDNA substrates were prepared by mixing ssDNA oligos to a final concentration of 80 μM in annealing buffer (20 mM HEPES pH7.5, 200 mM KCl), denaturation at 95 °C for 10 min and gradually temperature decrease to 4 °C during 20 minutes in a thermal cycler (Applied Biosystems). Ribonucleoprotein complexes (RNP) of CasΦ3 were formed by mixing an equal volume of 50 μM CasΦ3 and 50 μM CasΦ3 mature crRNA (IDT).

For specific dsDNA cleavage assays, FAM-labeled dsDNA substrates were incubated at 400 nM with 2 μM of CasΦ3 RNP in cleavage buffer (20 mM HEPES pH7.5, 160 mM KCl, 10% glycerol, 5 mM MgCl_2_) for 2h at 37 °C, or as otherwise stated in the figure legends. For ion dependency assays (Extended Data Fig. 3e) 5mM MgCl_2_ was substituted by 5mM Ethylenediaminetetraacetic acid (EDTA), CaCl_2_, MnCl_2_, FeSO_4_, CoCl_2_, NiSO_4_, CuCl_2_, ZnSO_4_. For DNA saturation experiments (Extended Data Fig. 3f) 1uM of CasΦ3 RNP was incubated with 0.5−8 uM of labelled dsDNA for 2h at 37°C. For non-specific trans ssDNA cleavage assays (Extended Data Fig. 2b-c, 8b-c), 0.4 μM FAM-labeled non-specific ssDNA substrate (i.e., not complementary to the crRNA) was incubated with 2 μM CasΦ3 RNP as described above, along with 0.1 μM of unlabeled activator ssDNA or dsDNA (complementary to the crRNA) in cleavage buffer for 1 h at 37°C. The reactions were stopped by adding equal volumes of stop buffer (8 M Urea, 100 mM EDTA at pH8) followed by incubation at 95°C for 5 min. Cleavage products were resolved on 15% Novex TBE-Urea Gels (Invitrogen), run according to manufacturer’s instructions. Gels were imaged using an Odyssey FC Imaging System (Li-Cor). Densitometric analysis of bands in gels was performed using ImageJ. The cleavage efficiency was calculated as the intensity of the bands corresponding to the products divided by the total intensity for the specific dsDNA cleavage assays, or as the depletion of signal of the non-cleaved product for non-specific ssDNA degradation assays.

### Sample preparation for Cryo-EM

For the preparation of the Cryo-EM sample, Ni^2+^ was used as a catalytic ion instead of Mg^2+^ due to the higher yield obtained with this metal. CasΦ3 RNP was prepared as described before. 25 nmol of RNP and 37 nmol of unlabeled dsDNA substrate were incubated in 25 ml of MonoQ A buffer (20 mM HEPES pH7.5, 200 mM KCl, 1 mM NiSO_4_, 0.5 mM TCEP) for 2h at 20°C to allow DNA cleavage. The product of the reaction was loaded in a MonoQ column equilibrated with MonoQ A buffer, and CasΦ3 R-loop complex was separated from the RNP and the unbound DNA substrate by a salt gradient elution using MonoQ B buffer (20 mM HEPES pH7.5, 2 M KCl, 1 mM NiSO_4_, 0.5 mM TCEP). CasΦ3 R-loop eluted at 16-20 % of MonoQ buffer B (~500 mM KCl). The R-loop complex was further purified from unbound DNA by SEC using a Superdex 200 Increase 10/300 GL column (Cytiva) equilibrated with MonoQ A buffer. The molecular weight of the complex and the sample homogeneity was estimated using a Refeyn One mass photometer (Refeyn), using 10-20 nM of protein diluted in MonoQ A buffer (Extended Data Fig. 3A). 2.5 μL of freshly purified CasΦ3 R-loop complex (Absorbance_260 nm_ of ~1.6) was applied to UltrAuFoil 300 mesh R0.6/1.0 holey grids (Quantifoil), glow-discharged for 60 s at 10 mA (Leica EM ACE200), and plunge-frozen in liquid ethane (pre-cooled with liquid nitrogen) using a Vitrobot Mark IV (FEI, Thermo Fisher Scientific) using the next conditions: blotting time 3 s, 100% humidity and 4° C.

### CryoEM Data Collection and Processing

Movies were collected on Titan Krios G3 Cryo-TEM equipped with a TFS Falcon III camera operated at 300 keV in counting mode. Exposure 1.05 e/Å^2^/frame, in 40 frames and hence a final dose of 42 e/Å^2^. The calibrated pixel size was 0.832 Å/px. All movies were pre-processed using WARP 1.0.9^31^ (Extended Data Fig. 3). Motion correction was performed with a temporal resolution of 20 for the global motion and 5◻×◻5 spatial resolution for the local motion. We considered motion in the 45–3 Å range weighted with a B-factor of −500◻Å^2^. Only Micrographs displaying less than 5 Å intraframe motion were used. CTF estimation was performed using 5◻×◻5 patches in the 35-4 Å range. We selected micrographs with fitted defocus between 0.0 and 5.0◻μm, and a resolution better than 5◻Å. For the particle picking, the micrographs were masked, and particles were picked using a re-trained BoxNet deep convolutional neural network. This resulted in 3,504,102 particles from 4,393 micrographs. Particles were extracted with a box size of 256×256 and a pixel size of 0.832 which were inverted and normalized before being imported into RELION 3.1 ^32^ for 2D classification. The selected 2D classes were imported in cryoSPARC 3.1.0 ^33^ where they were 3D classified into four initial classes. The volume with the largest number of particles was 3D autorefined to an initial 2.61 Å resolution map. The conformational heterogeneity of the particles used in this volume was inspected through a 3D variability analysis job, and the two more divergent volumes were used as input for heterogeneous refinement. The 3D variability of the particles in the best volume was further analysed followed by heterogeneous refinement with four classes. The resulting four volumes were non-uniform refined to obtain maps at 2.7−3.3 Å resolution. The two best maps (2.7 and 2.9 Å resolution) represent the different conformational states of the complex that are discussed in the text. Sharpened and local resolution maps were calculated with PHENIX^34^, and directional resolution anisotropy analysis were performed with the 3D-FSC server ^35^.

### Atomic model building and refinement

An initial model containing the complete DNA and RNA sequence and ~50% of the protein sequence was built ab initio using map-to-model implemented in PHENIX^34^. COOT^36^ was used to connect, extend and correct the protein fragments to generate a model covering ~70% of the protein sequence. The rest of the model was autobuilt by using buccaneer implemented in CCP-EM^37^, and subsequently corrected in COOT. The final model was obtained after several rounds of refinement using phenix.real_space_refine and manual inspection and correction in COOT. The final model covers 92% of the protein sequence, mainly lacking a C-terminal segment predicted to be unstructured. Map and molecular model images were created using ChimeraX38.

## Data Availability

The atomic coordinates and cryo-EM maps have been deposited in the Protein Data Bank and EMDB under accession codes 7ODF and EMD-12827. All other data are available from the corresponding author upon request.

## Extended Data Figure Legends

**Extended Data Figure 1.- Sequence alignment of known members of the CasΦ family**. The amino acid sequences of CasΦ 1 to 10 were aligned by Clustal Omega (http://www.ebi.ac.uk/Tools/msa/clustalo). The figure was prepared with ESPript (http://espript.ibcp.fr). Residue numbers are labelled according to the CasΦ3 sequence. Similar residues are shown in red and identical residues in white over red background. The different structural domains of CasΦ3 and their amino acid composition are depicted as boxes above the sequences and labelled with the same names and colour code as in Fig. 1a. The mutations performed in key residues commented through the text are marked below.

**Extended Data Figure 2.- Cas**Φ3 **endonuclease biochemical characterisation. a)** representative dsDNA cleavage pattern generated by CasΦ3 wild type (WT). T-strand (TS) and NT-strand (NTS) products are marked, showing a cut at position −13, −14 and −15 of the NT-strand, while the T-strand is cleaved at position +23. The sequence of the double labeled duplex is shown below, marking the position of the cut (triangles), and the size of the labelled products. **b)** Unspecific ssDNA degradation after activation with a specific target ssDNA of different length (Oligonucleotides T-AAG-3 to T-AAG-30 in Extended Data Table II). **c)** Unspecific ssDNA degradation after activation with a specific dsDNA activator of different lengths (Oligonucleotides T-AAG-3/NT-TTC-3 to T-AAG-30/NT-TTC-30 in Extended Data Table II). **d)** Schematic cartoon of the results shown in b) and c). Activation of the unspecific ssDNA cleavage is observed between 12-30 nt. (i) The RuvC domain of CasΦ3 RNP is inhibited. Full activation of the unspecific cleavage is observed when using a ssDNA or dsDNA activator pairing with the crRNA between 12-18 nt (ii and iv). The use of longer oligos as ssDNA(iii) or dsDNA (v) result in a reduction of the cleavage efficiency, likely due to a steric occlusion of the catalytic site by the T-strand and NT-strand. **e)** DNA cleavage dependency on divalent metal ions. Mg^2+^, Mn^2+^, Fe^2+^, Co^2+^ and Ni^2+^ metal ions support CasΦ3 catalytic activity, while Ca^2+^, Cu^2+^, Zn^2+^ do not. Depletion of the cation by EDTA abrogates phosphodiester hydrolysis. **f)** Cleavage assay using the target dsDNA shows the cleavage products of the different strands at different enzyme and substrates ratios. Quantification of the cleaved and non-cleaved dsDNA substrate is shown in the chart as mean ± s.d. (see Methods). The curve shows an increase of the non-cleaved substrate when a 1:1 ratio is reached. An asymptotic behaviour is observed for the NT-strand products. **g)** Time course of the cleavage reaction by CasΦ3. CasΦ3 endonuclease completes the reaction in approximately 120◻min for the T-strand while the NT-strand cleavage is completed in 20 min. **h)** Time course of the cleavage reaction by CasΦ3-ΔCT mutant lacking the C-terminal 39 residues. Oligonucleotides 3F-T-AAG-30/NT-TTC-30, T-AAG-30/5F-NT-TTC-30 and 3F-T-AAG-30/5F-NT-TTC-30 (Extended Data Table II) were used as substrate to visualize the cleavage of the target, non-target and both DNA strands in a) d-g). Experiments displayed are representative of at least three replicates.

**Extended Data Figure 3.- Single particle cryo-EM analysis of the Cas**Φ3 **endonuclease R-loop complex. a)** SDS-PAGE, electrophoretic mobility shift assay (EMSA), and denaturing polyacrylamide gels showing the reconstitution of CasΦ3/R-loop complex. The complex shown in a) was reconstituted with the labeled oligonucleotides 3F-T-AAG-30/5F-NT-TTC-30 under similar conditions as the complex used for cryo-EM. The right-side panel shows the experimental mass and homogeneity of the complex used for cryo-EM (i.e. using the unlabeled nucleotides T-AAG-30/NT-TTC-30), estimated by mass photometry. **b)** Representative cryo-EM micrograph of the CasΦ3/R-loop complex in vitreous ice on gold grids before and after denoising with WARP^31^ **c)** Representative reference-free 2D class averages sorted by class distribution (RELION ^32^). **d)** Overview of the cryo-EM data processing workflow for the CasΦ3/R-loop complex.

**Extended Data Figure 4.- Resolution assessment, validation of cryoEM density maps. a, d)** Local resolution cryo-EM maps of the CasΦ3-R-loop complex. **b, e)** Angular distribution plot showing the relative orientation of particles in the final 3D reconstruction. **c, f)** Fourier shell correlation (FSC) curves and sphericity of the final cryo-EM density maps (3D-FSC ^35^, Extended Data Table I).

**Extended Data Figure 5.- Cas**Φ3**/R-loop atomic model and cryo-EM maps a)** Comparison of the cryo-EM maps showing the high flexibility of the NPID, RuvC and STP domains. **b)** View of the cryo-EM map in the RBD domain (contour level 7.5), **c)** PAM binding region (contour level 7.0), **d)** unzipping cavity (contour level 7.5), **e)** RuvC insertion (contour level 7.8), **f)** hydrophobic plug (contour level 6) and **g)** the RuvC catalytic site (contour level 5.7).

**Extended Data Figure 6.- Protein-nucleic acids interactions in the Cas**Φ3**-R-loop structure.** Polar and non-polar contacts of the nucleic acids with the protein side and main chain are indicated (see key in the figure).

**Extended Data Figure 7.- The cleaved NT-strand of the Cas**Φ3**-R-loop complex. a-b)** Two different views of the cryo-EM map density observed at low contour suggesting that the dinucleotide in the catalytic site belongs to the NT-strand. The nucleotides in blue are shown in a) for illustrative purposes only. **c)** EMSA assay using FAM labelled NT- and T-strand showing that the PAM distal region of the substrate dissociates of the R-loop complex after cleavage supporting that the dinucleotide in the catalytic site belongs to the NT-strand.

**Extended Data Figure 8.- PAM specificity and crRNA/DNA hybrid assembly. a)** cleavage assay with CasΦ3 WT and PAM interacting mutants, using target dsDNA as substrate containing different PAM or no PAM sequence. Oligonucleotides 3F-T-AAG-30/5F-NT-TTC-30, 3F-T-TTG-30/5F-NT-AAC-30, 3F-T-CCG-30/5F-NT-GGC-30, 3F-T-GGG-30/5F-NT-CCC-30 and 3F-T-30/5F-NT-30 were used as activators (Extended Data Table II) **b)** CasΦ3 activation of unspecific ssDNA degradation assay using an 18-nt dsDNA containing different PAM or no PAM sequence as activator. **c)** Unspecific ssDNA degradation by CasΦ3 WT and representative mutants involved in the PAM recognition (K30A/Q123A/Q197A), unwinding (K55A), and RuvC insertion (R643A) after activation with a 18-nt ssDNA without the PAM or a 18-nt dsDNA with the PAM. **d**) schematic representation explaining the results of the experiments shown in c). Gels shown are representative of three independent experiments.

**Extended Data Figure 9.- Structural comparison of Cas**Φ3 **RuvC domain with other Cas nucleases.** A structural homology search of CasΦ3 against the PDB was performed using DALI ^39^. Only the RuvC domain displays homology with other Cas nucleases. **a)** Superposition of CasΦ3 and Cas12f. **b)** Superposition of CasΦ3 and Cas12g. Both Cas12f and Cas12g present a Zn^2+^ atom coordinated by 4 conserved cysteines as CasΦ3. The rest of the domain is different to the TNB domain in Cas12f and Cas12g. **c)** Superposition of CasΦ3 and Cas12b. **d)** Superposition of CasΦ3 and Cas12i. Both Cas12b and Cas12i present DNA in the catalytic site and the presence of the Nuc domain inserted in the RuvC.

## Extended Data Tables

**Extended Data Table I.** Cryo-EM data collection, refinement and validation statistics

|  | (EMDB-12827)<br>(PDB 7ODF) | (EMDB-Waiting<br>for code) |
| --- | --- | --- |
| **Data collection and processing** |  |  |
| Voltage (kV) | 300 | 300 |
| Total electron exposure (e-/Å<sup>2</sup>) | 42 (1.05/frame) | 42 (1.05/frame) |
| Defocus range (μm) | 0.8/2.6 | 0.8/2.6 |
| Pixel size (Å) | 0.832 | 0.832 |
| Symmetry imposed | n/a | n/a |
| Initial particle images (no.) | 3,504,102 | 3,504,102 |
| Final particle images (no.) | 178,412 | 110,551 |
| Sphericity | 0.955 | 0.856 |
| Map resolution (Å) | 2.66 | 2.84 |
| FSC threshold | 0.143 | 0.143 |
| Map resolution range (Å) | 2.66-10.19 | 2.87-11.13 |
| **Refinement** |  |  |
| Initial model used (PDB code) | none |  |
| Model resolution (Å) | 2.92 |  |
| FSC threshold | 0.143 |  |
| Model resolution range (Å) | 2.7-7.3 |  |
| Map sharpening *B* factor (Å<sup>2</sup>) | -67.26 |  |
| Model composition |  |  |
| Non-hydrogen atoms | 6852 |  |
| Protein residues | 703 |  |
| Nucleotides | 78 |  |
| Ligands | Ni:1 Zn:1 |  |
| Water | 5 |  |
| *B* factors (Å<sup>2</sup>) | Min/max/mean |  |
| Protein | 51.88/144.89/80.88 |  |
| Nucleotide | 56.10/142.61/79.83 |  |
| Ligand | 99.25/127.66/113.45 |  |
| Water | 80.64/80.64/80.64 |  |
| R.m.s. deviations |  |  |
| Bond lengths (Å) | 0.004 (0) |  |
| Bond angles (°) | 0.825 (0) |  |
| Validation |  |  |
| MolProbity score | 1.41 |  |
| Clashscore | 7.53 |  |
| Rotamer outliers (%) | 0.78 |  |
| CaBLAM outliers (%) | 0.72 |  |
| Cβ outliers (%) | 0.0 |  |
| Peptide plane (%) |  |  |
| Cis proline/general | 0.0/0.0 |  |
| Twisted proline/general | 0.0/0.0 |  |
| Ramachandran plot |  |  |
| Favored (%) | 98.43 |  |
| Allowed (%) | 1.43 |  |
| Disallowed (%) | 0.0 |  |

**Extended Data Table II.**
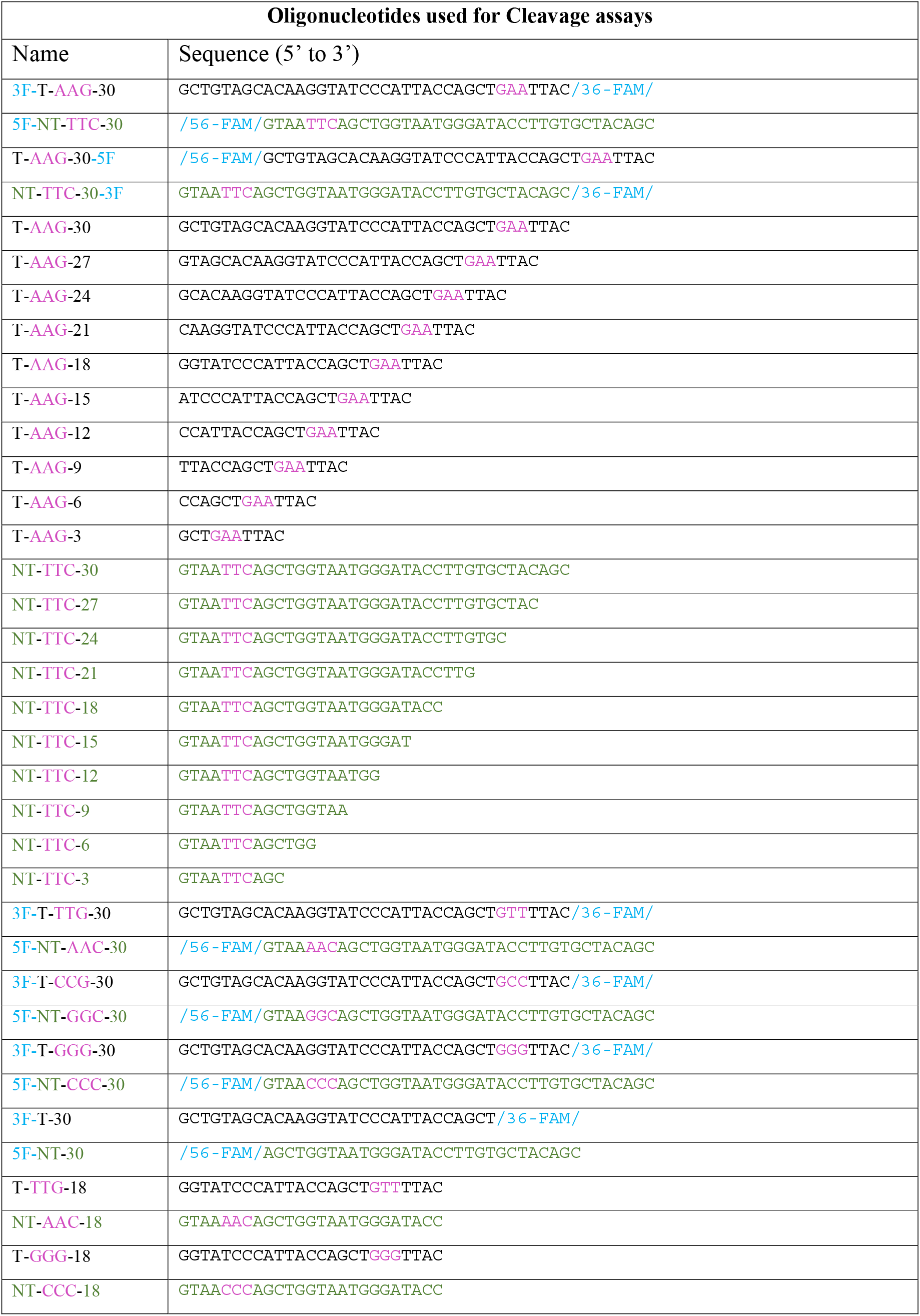
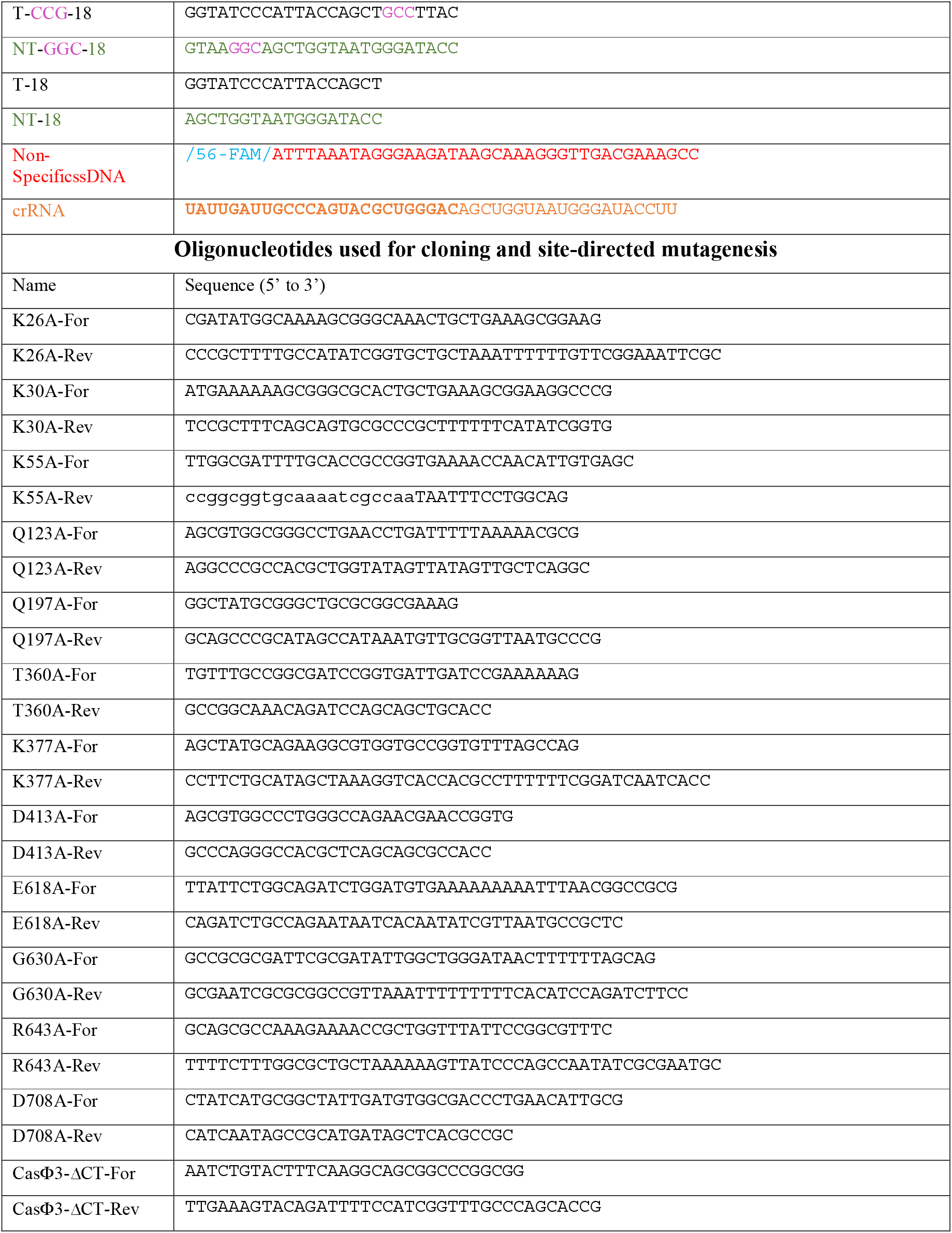
Oligonucleotides and primer sequences used in this study

**Supplementary Video 1.**
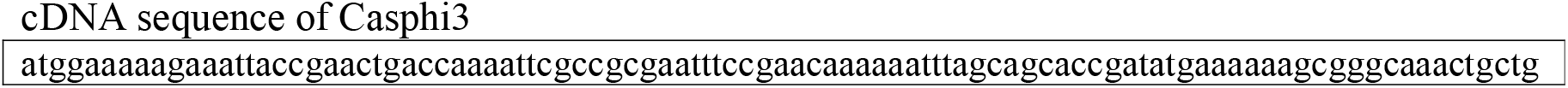
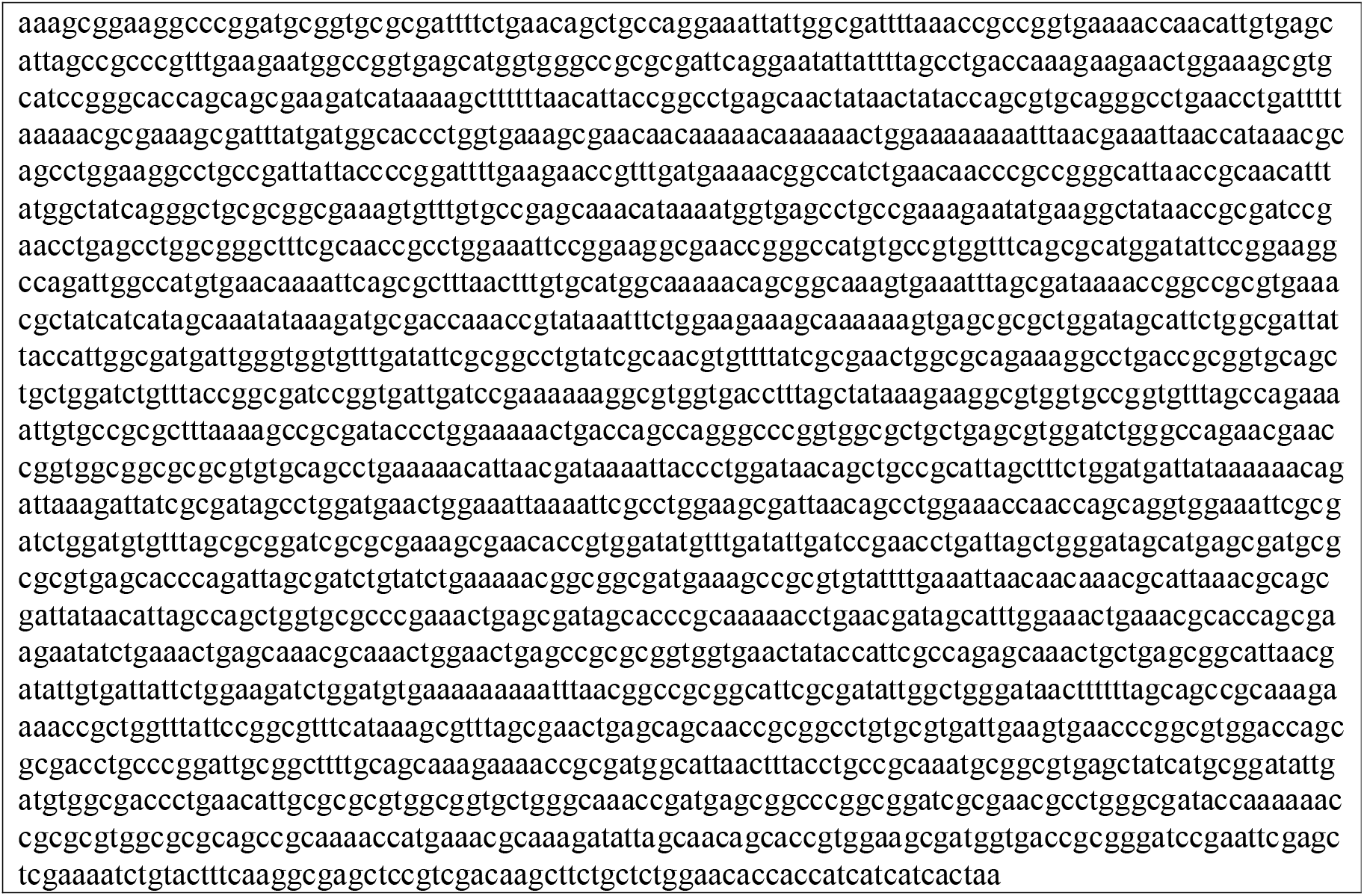
Conformational heterogeneity of the particles. The video displays the 3D variability analysis performed with cryoSPARC. Large variability is observed in the NPID, STP and RuvC domains.

## References

1 Al-Shayeb, B. et al. Clades of huge phages from across Earth’s ecosystems. Nature 578, 425–431, doi:10.1038/s41586-020-2007-4 (2020).

2 Pausch, P. et al. CRISPR-CasPhi from huge phages is a hypercompact genome editor. Science 369, 333–337, doi:10.1126/science.abb1400 (2020).

3 Abudayyeh, O. O. et al. C2c2 is a single-component programmable RNA-guided RNA-targeting CRISPR effector. Science 353, aaf5573, doi:10.1126/science.aaf5573 (2016).

4 Horvath, P. & Barrangou, R. CRISPR/Cas, the immune system of bacteria and archaea. Science 327, 167–170, doi:10.1126/science.1179555 (2010).

5 Sorek, R., Lawrence, C. M. & Wiedenheft, B. CRISPR-mediated adaptive immune systems in bacteria and archaea. Annu Rev Biochem 82, 237–266, doi:10.1146/annurev-biochem-072911-172315 (2013).

6 Adli, M. The CRISPR tool kit for genome editing and beyond. Nat Commun 9, 1911, doi:10.1038/s41467-018-04252-2 (2018).

7 Hsu, P. D., Lander, E. S. & Zhang, F. Development and applications of CRISPR-Cas9 for genome engineering. Cell 157, 1262–1278, doi:10.1016/j.cell.2014.05.010 (2014).

8 Cox, D. B. T. et al. RNA editing with CRISPR-Cas13. Science 358, 1019–1027, doi:10.1126/science.aaq0180 (2017).

9 Walton, R. T., Christie, K. A., Whittaker, M. N. & Kleinstiver, B. P. Unconstrained genome targeting with near-PAMless engineered CRISPR-Cas9 variants. Science 368, 290–296, doi:10.1126/science.aba8853 (2020).

10 Barrangou, R. & Horvath, P. A decade of discovery: CRISPR functions and applications. Nat Microbiol 2, 17092, doi:10.1038/nmicrobiol.2017.92 (2017).

11 Saha, K. et al. The NIH Somatic Cell Genome Editing program. Nature 592, 195–204, doi:10.1038/s41586-021-03191-1 (2021).

12 Chen, J. S. et al. CRISPR-Cas12a target binding unleashes indiscriminate single-stranded DNase activity. Science 360, 436–439, doi:10.1126/science.aar6245 (2018).

13 Stella, S., Alcon, P. & Montoya, G. Structure of the Cpf1 endonuclease R-loop complex after target DNA cleavage. Nature 546, 559–563, doi:10.1038/nature22398 (2017).

14 Sternberg, S. H., Redding, S., Jinek, M., Greene, E. C. & Doudna, J. A. DNA interrogation by the CRISPR RNA-guided endonuclease Cas9. Nature 507, 62–67, doi:10.1038/nature13011 (2014).

15 Jiang, F. et al. Structures of a CRISPR-Cas9 R-loop complex primed for DNA cleavage. Science 351, 867–871, doi:10.1126/science.aad8282 (2016).

16 Anders, C., Niewoehner, O., Duerst, A. & Jinek, M. Structural basis of PAM-dependent target DNA recognition by the Cas9 endonuclease. Nature 513, 569–573, doi:10.1038/nature13579 (2014).

17 Jiang, F. & Doudna, J. A. CRISPR–Cas9 Structures and Mechanisms. Annual Review of Biophysics 46, 505–529, doi:10.1146/annurev-biophys-062215-010822 (2017).

18 Stella, S., Alcon, P. & Montoya, G. Class 2 CRISPR-Cas RNA-guided endonucleases: Swiss Army knives of genome editing. Nat Struct Mol Biol 24, 882–892, doi:10.1038/nsmb.3486 (2017).

19 Jiang, F., Zhou, K., Ma, L., Gressel, S. & Doudna, J. A. STRUCTURAL BIOLOGY. A Cas9-guide RNA complex preorganized for target DNA recognition. Science 348, 1477–1481, doi:10.1126/science.aab1452 (2015).

20 Stella, S. et al. Conformational Activation Promotes CRISPR-Cas12a Catalysis and Resetting of the Endonuclease Activity. Cell 175, 1856–1871 e1821, doi:10.1016/j.cell.2018.10.045 (2018).

21 Swarts, D. C. & Jinek, M. Mechanistic Insights into the cis- and trans-Acting DNase Activities of Cas12a. Mol Cell 73, 589–600 e584, doi:10.1016/j.molcel.2018.11.021 (2019).

22 Swarts, D. C., van der Oost, J. & Jinek, M. Structural Basis for Guide RNA Processing and Seed-Dependent DNA Targeting by CRISPR-Cas12a. Mol Cell 66, 221–233 e224, doi:10.1016/j.molcel.2017.03.016 (2017).

23 Yamano, T. et al. Crystal Structure of Cpf1 in Complex with Guide RNA and Target DNA. Cell 165, 949–962, doi:10.1016/j.cell.2016.04.003 (2016).

24 Takeda, S. N. et al. Structure of the miniature type V-F CRISPR-Cas effector enzyme. Mol Cell 81, 558–570.e553, doi:10.1016/j.molcel.2020.11.035 (2021).

25 Li, Z., Zhang, H., Xiao, R., Han, R. & Chang, L. Cryo-EM structure of the RNA-guided ribonuclease Cas12g. Nature Chemical Biology 17, 387–393, doi:10.1038/s41589-020-00721-2 (2021).

26 Nowotny, M. Retroviral integrase superfamily: the structural perspective. EMBO Rep 10, 144–151, doi:10.1038/embor.2008.256 (2009).

27 Steitz, T. A. & Steitz, J. A. A general two-metal-ion mechanism for catalytic RNA. Proc Natl Acad Sci U S A 90, 6498–6502, doi:10.1073/pnas.90.14.6498 (1993).

28 Bartesaghi, A., Matthies, D., Banerjee, S., Merk, A. & Subramaniam, S. Structure of β-galactosidase at 3.2-Å resolution obtained by cryo-electron microscopy. Proceedings of the National Academy of Sciences 111, 11709, doi:10.1073/pnas.1402809111 (2014).

29 Chapados, B. R. et al. Structural biochemistry of a type 2 RNase H: RNA primer recognition and removal during DNA replication. J Mol Biol 307, 541–556, doi:10.1006/jmbi.2001.4494 (2001).

30 Kanaya, S. Enzymatic activity and protein stability of E. coli ribonuclease HI. Ribonucleases H., 1–38 (1998).

31 Tegunov, D. & Cramer, P. Real-time cryo-electron microscopy data preprocessing with Warp. Nature Methods 16, 1146–1152, doi:10.1038/s41592-019-0580-y (2019).

32 Zivanov, J. et al. New tools for automated high-resolution cryo-EM structure determination in RELION-3. Elife 7, doi:10.7554/eLife.42166 (2018).

33 Punjani, A., Rubinstein, J. L., Fleet, D. J. & Brubaker, M. A. cryoSPARC: algorithms for rapid unsupervised cryo-EM structure determination. Nature Methods 14, 290–296, doi:10.1038/nmeth.4169 (2017).

34 Liebschner, D. et al. Macromolecular structure determination using X-rays, neutrons and electrons: recent developments in Phenix. Acta Crystallogr D Struct Biol 75, 861–877, doi:10.1107/S2059798319011471 (2019).

35 Tan, Y. Z. et al. Addressing preferred specimen orientation in single-particle cryo-EM through tilting. Nat Methods 14, 793–796, doi:10.1038/nmeth.4347 (2017).

36 Emsley, P. & Cowtan, K. Coot: model-building tools for molecular graphics. Acta Crystallogr D Biol Crystallogr 60, 2126–2132, doi:10.1107/S0907444904019158 (2004).

37 Burnley, T., Palmer, C. M. & Winn, M. recent developments in the CCP-EM software suite. Acta Crystallographica Section D 73, 469–477, doi:doi:10.1107/S2059798317007859 (2017).

38 Goddard, T. D. et al. UCSF ChimeraX: Meeting modern challenges in visualization and analysis. Protein Sci 27, 14–25, doi:10.1002/pro.3235 (2018).

39 Holm, L. & Rosenström, P. Dali server: conservation mapping in 3D. Nucleic Acids Res 38, W545–549, doi:10.1093/nar/gkq366 (2010).

