## Supplementary material for "Structure of the mini-RNA-guided endonuclease CRISPR-CasΦ3": Supp fig 1

Extended Data Fig. 1

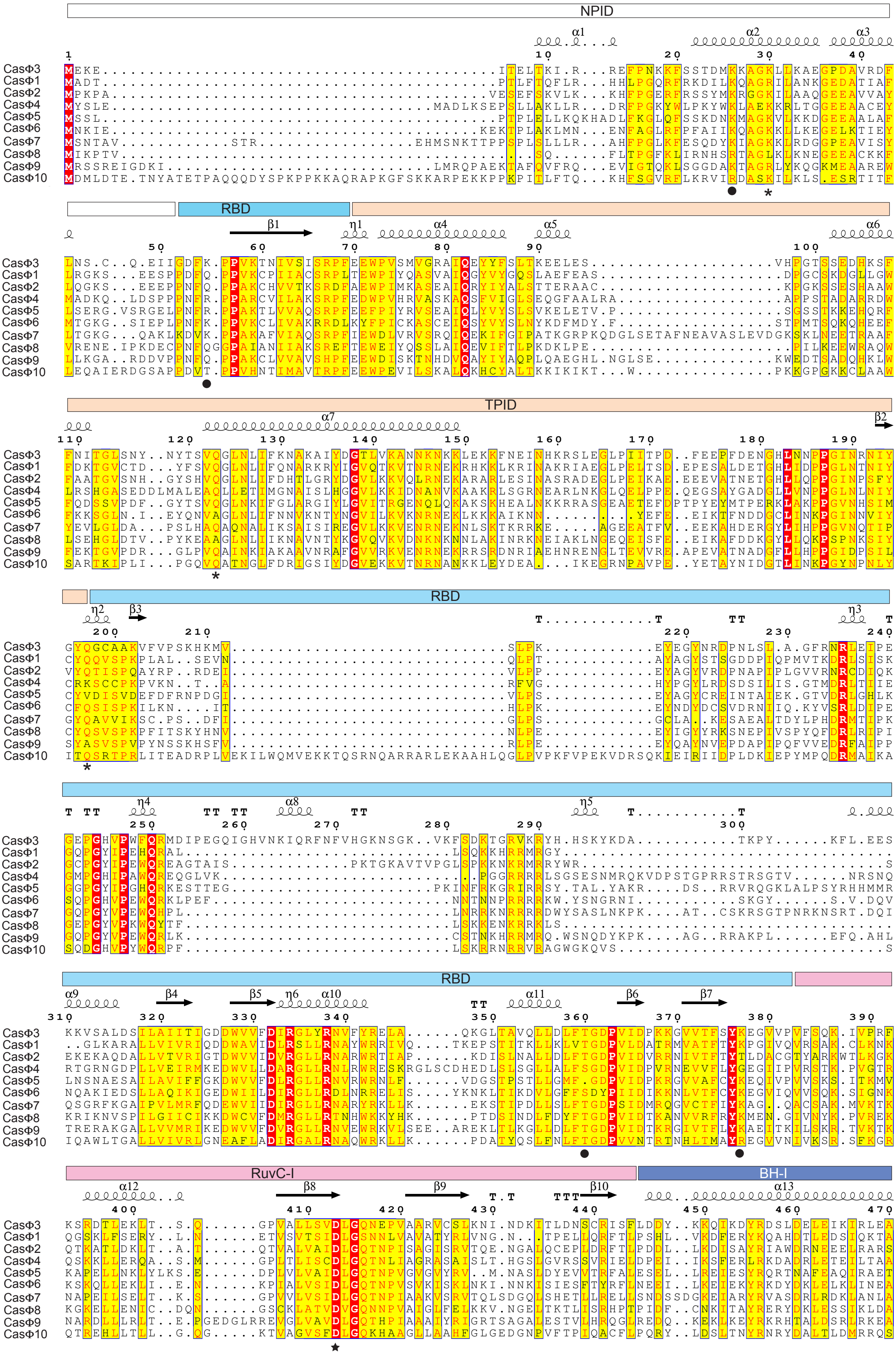


* PAM mutants
● Unwinding mutants
★ Catalytic mutants
■ RuvC insertion mutants
### STP and Plug mutants

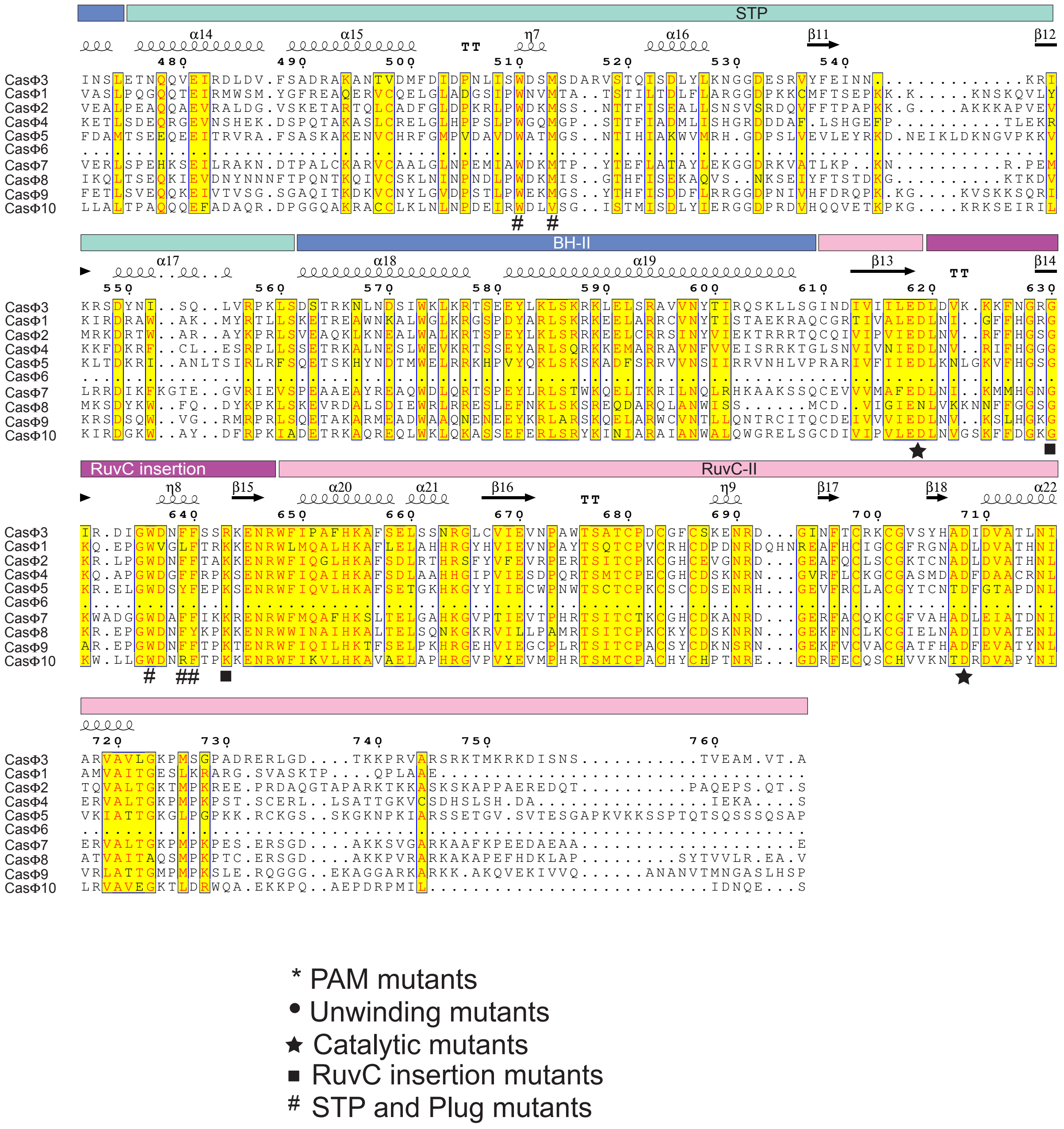
STP
BH-II
RuvC insertion
RuvC-II
CasΦ3
CasΦ1
CasΦ2
CasΦ4
CasΦ5
CasΦ6
CasΦ7
CasΦ8
CasΦ9
CasΦ10
* PAM mutants
● Unwinding mutants
★ Catalytic mutants
■ RuvC insertion mutants
### STP and Plug mutants
