## Supplementary figures and images for "Structure of the mini-RNA-guided endonuclease CRISPR-CasΦ3"

### Supp fig 2

#### Extended Data Fig. 2

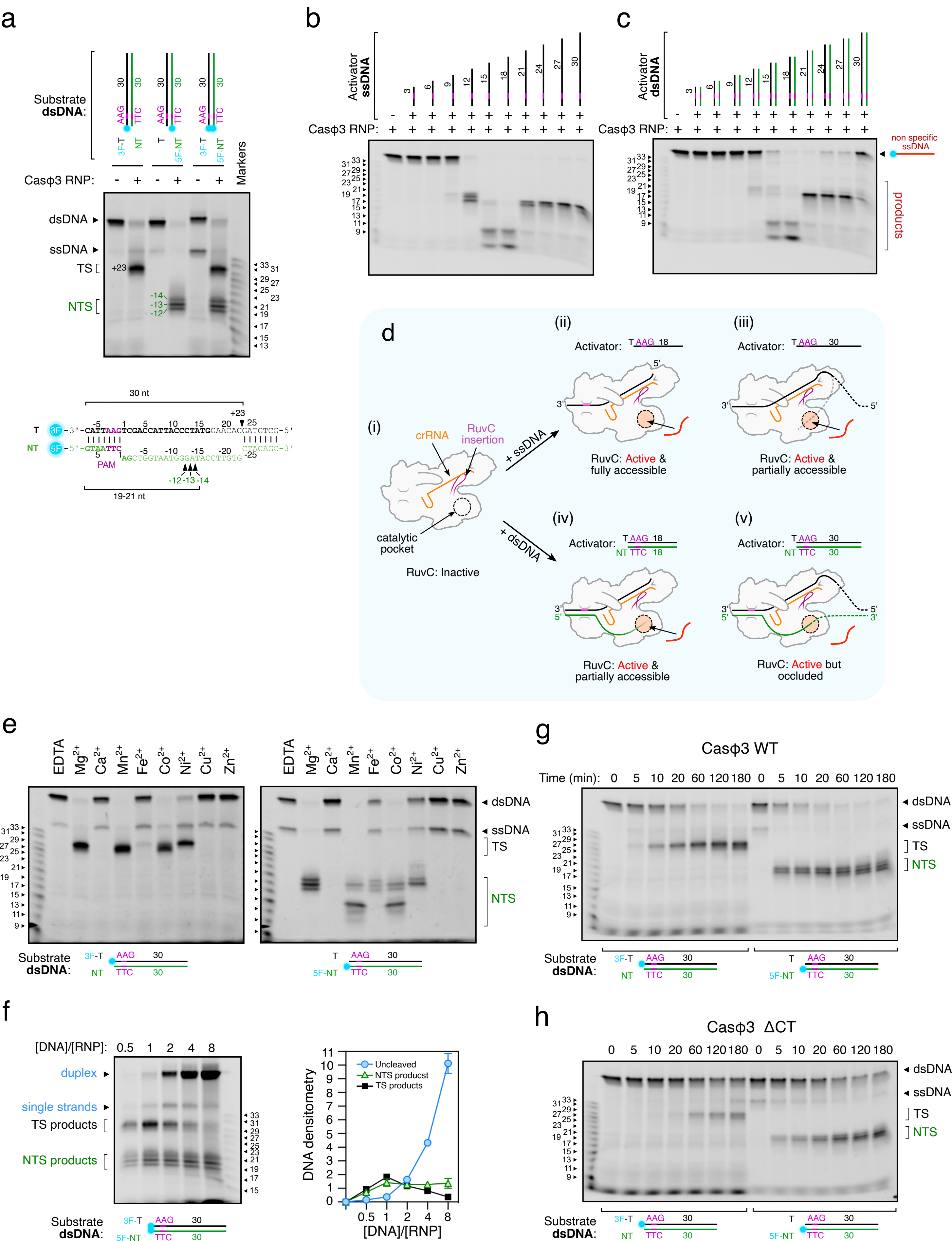

### Supp fig 4

Extended Data Fig. 4

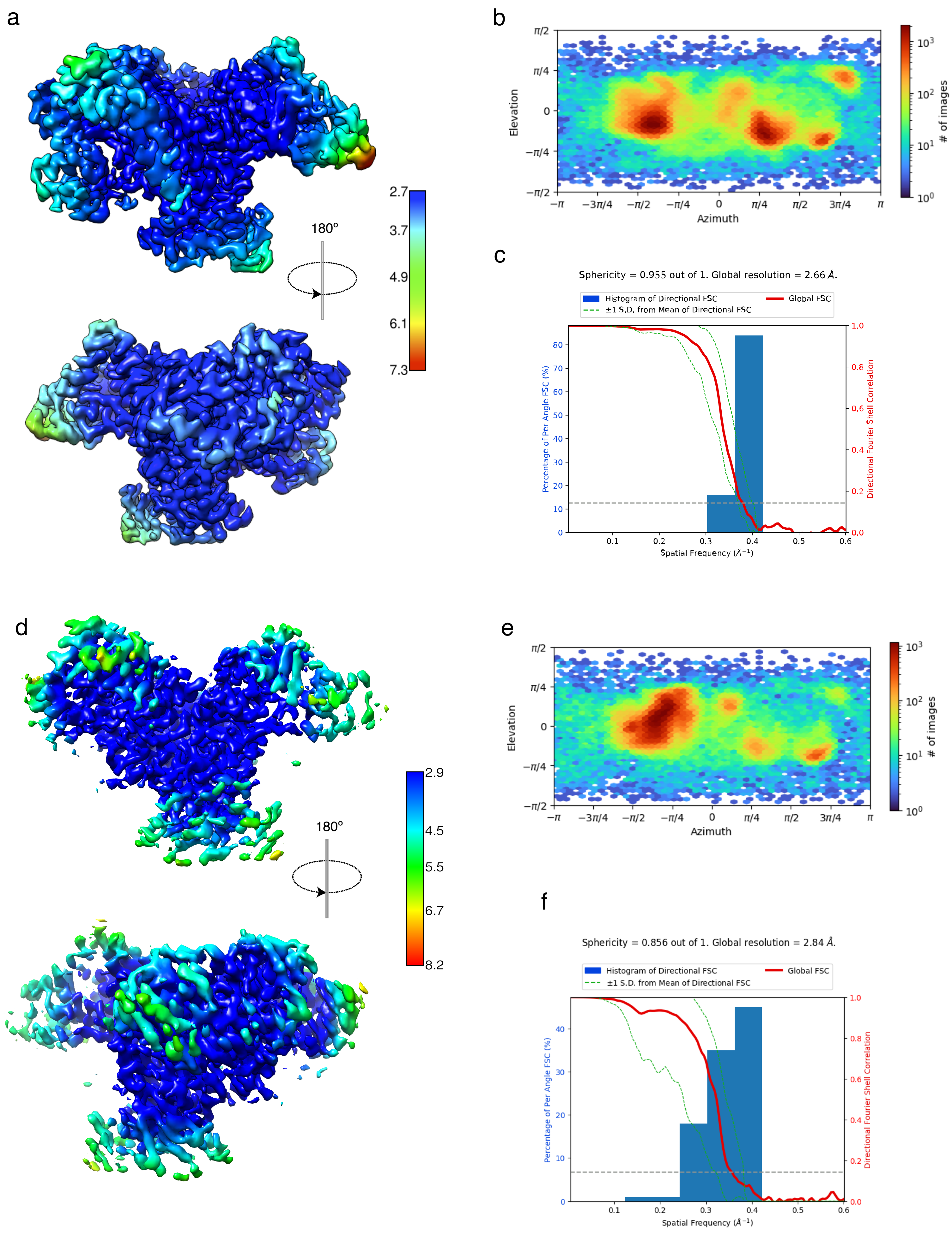

### Supp fig 5

Extended Data Fig. 5

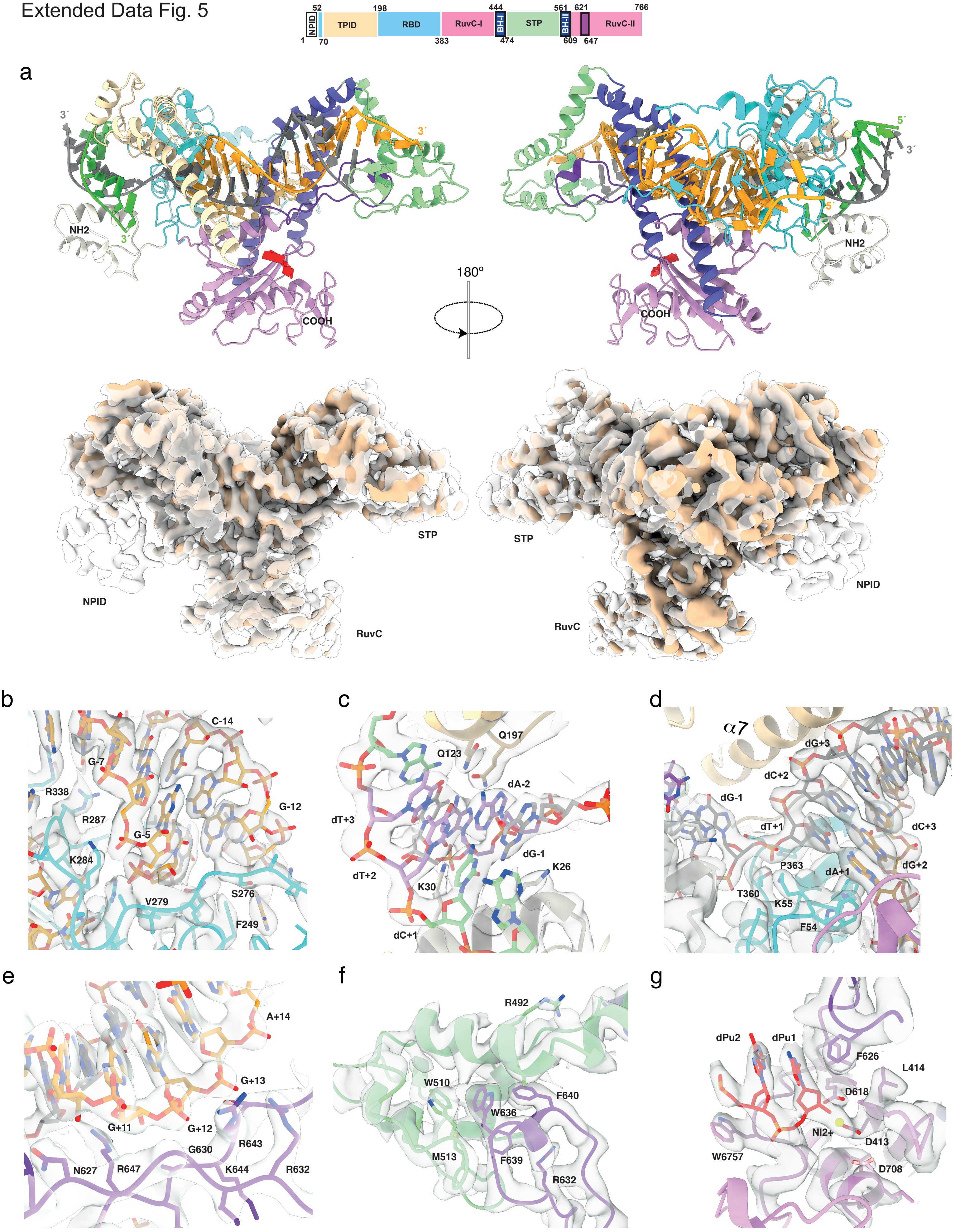

### Supp fig 6

#### Extended Data Fig. 6

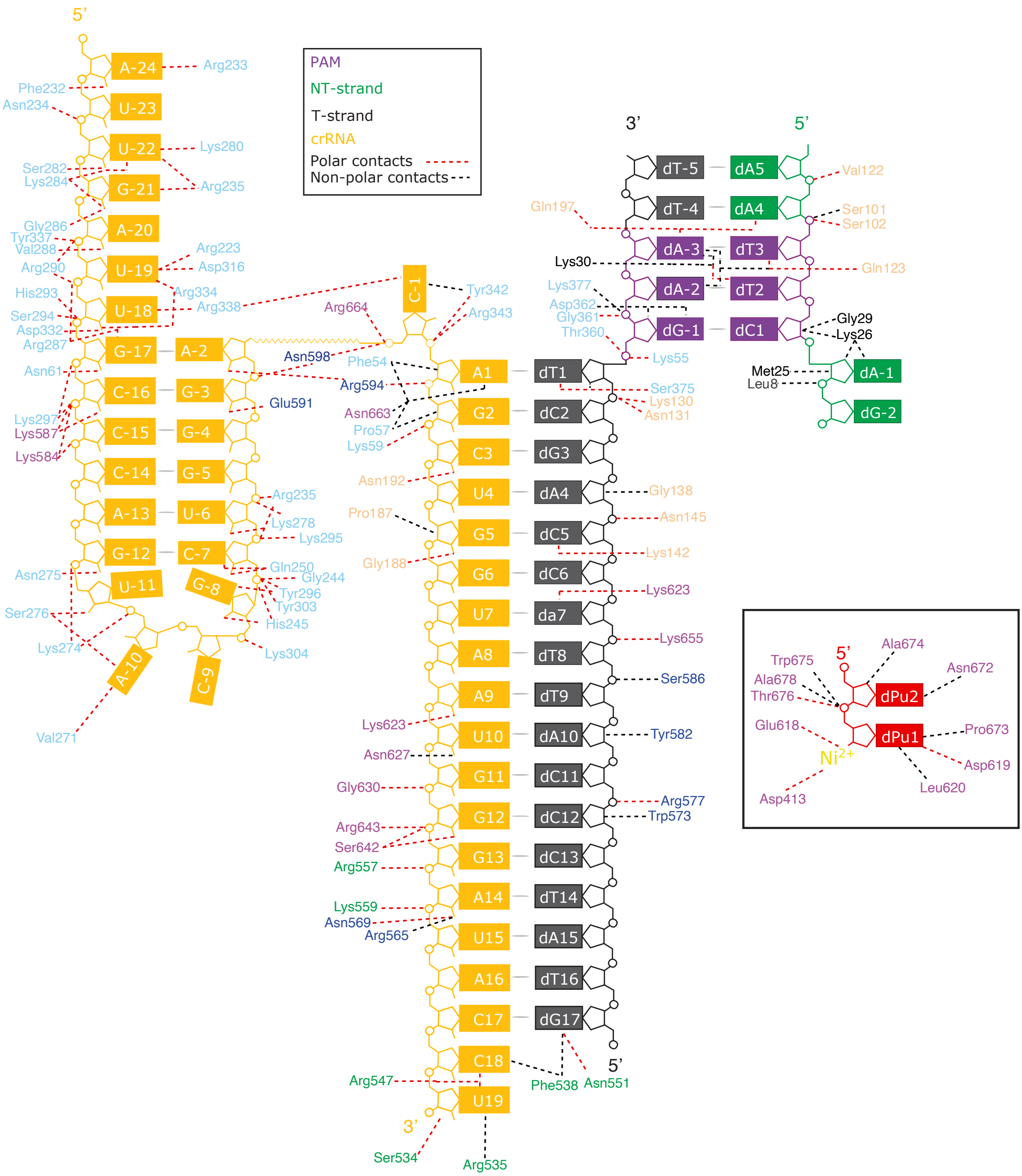

### Supp fig 7

Extended Data Fig. 7

a

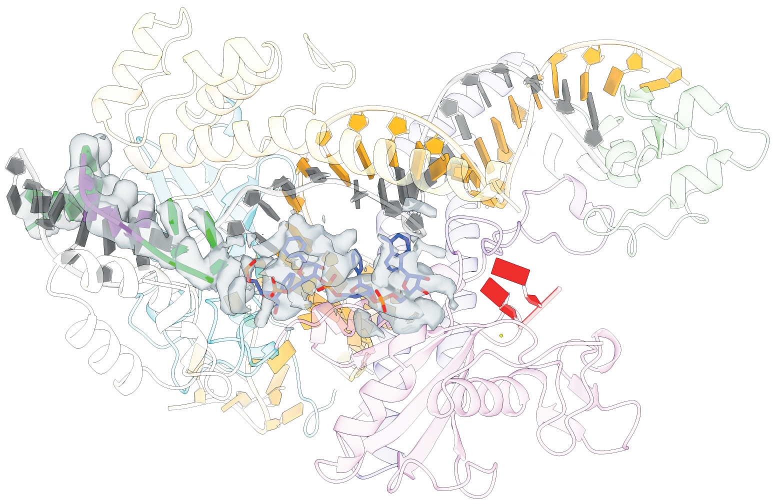

b

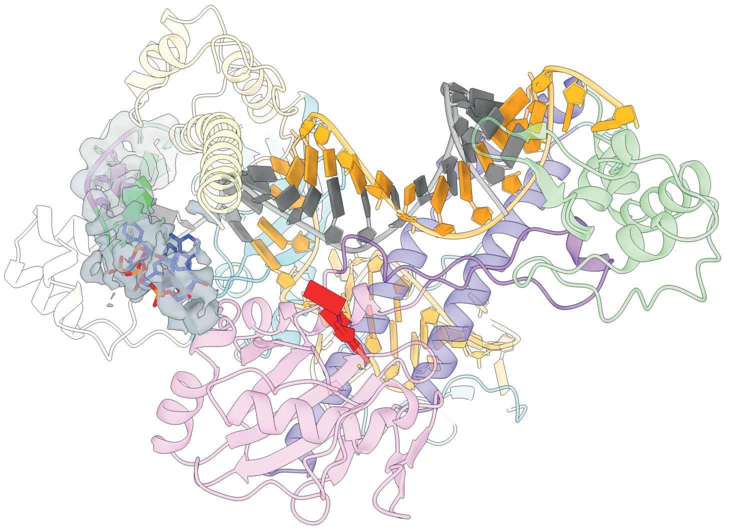

c

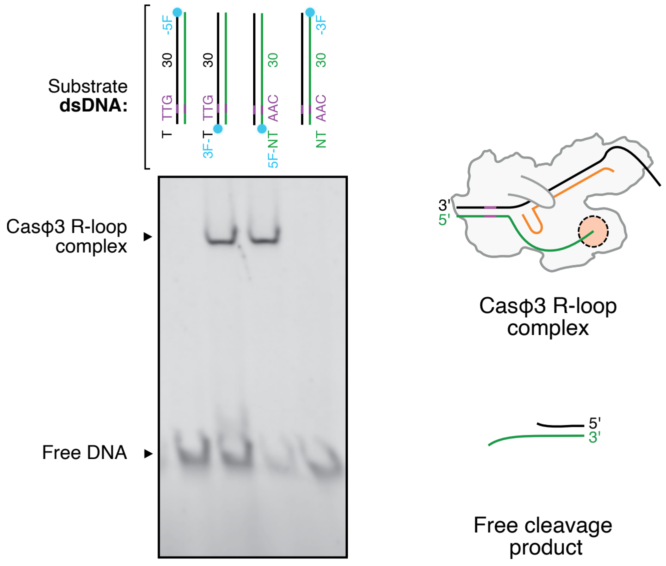

### Supp fig 8

#### Extended Data Fig. 8.

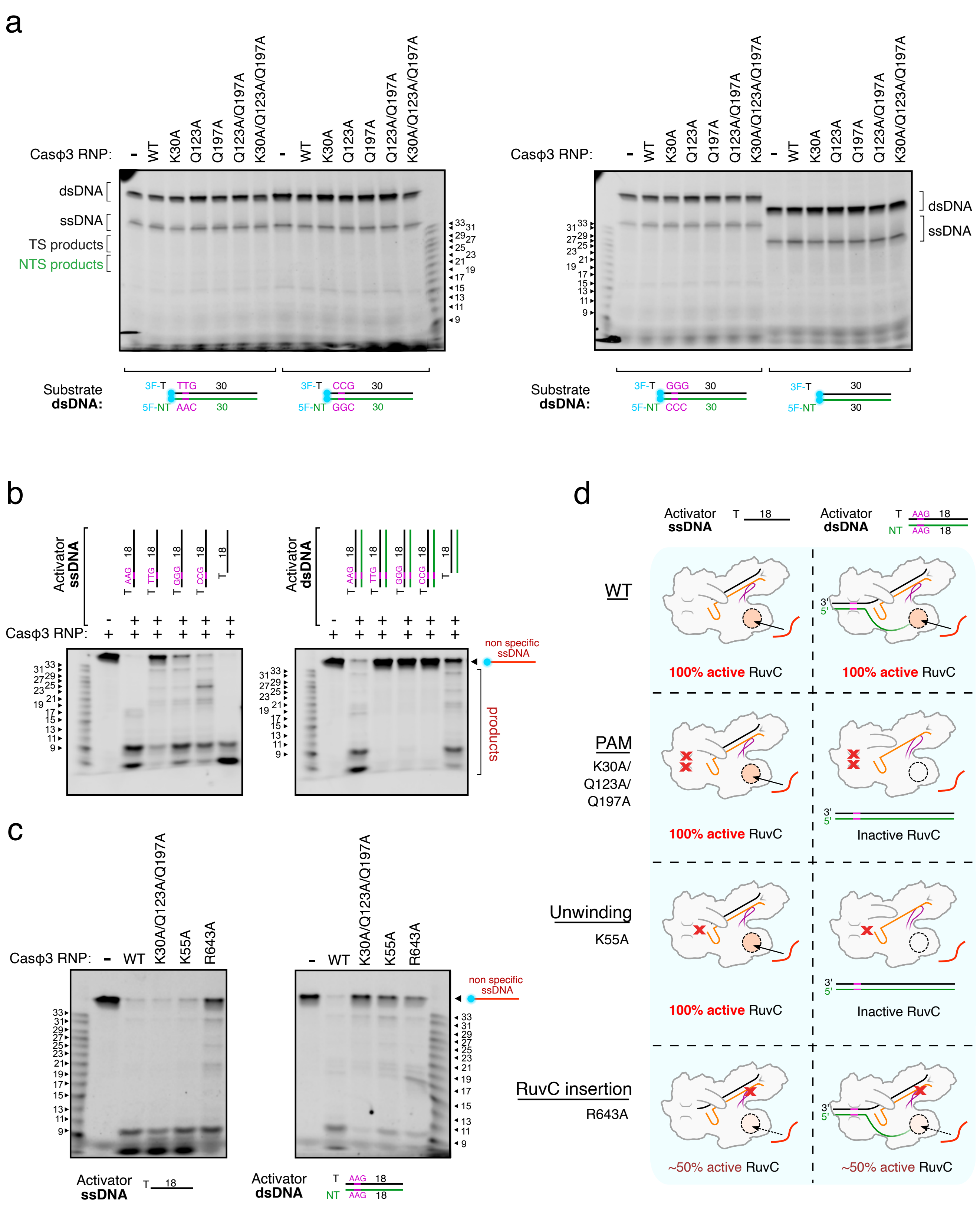

### Supp fig 9

Extended Data Fig. 9

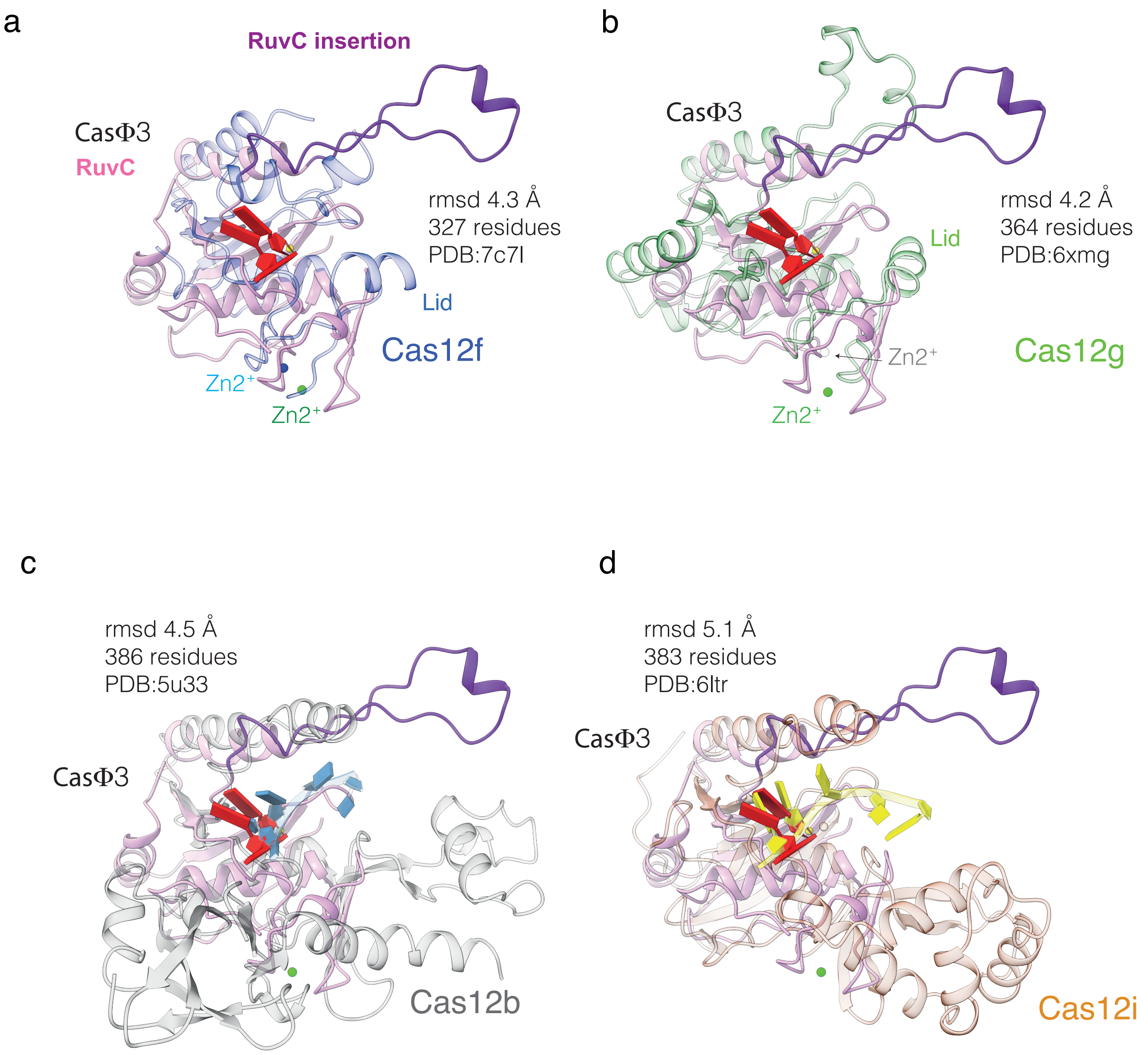

### Supp. fig 3

#### Extended Data Fig. 3

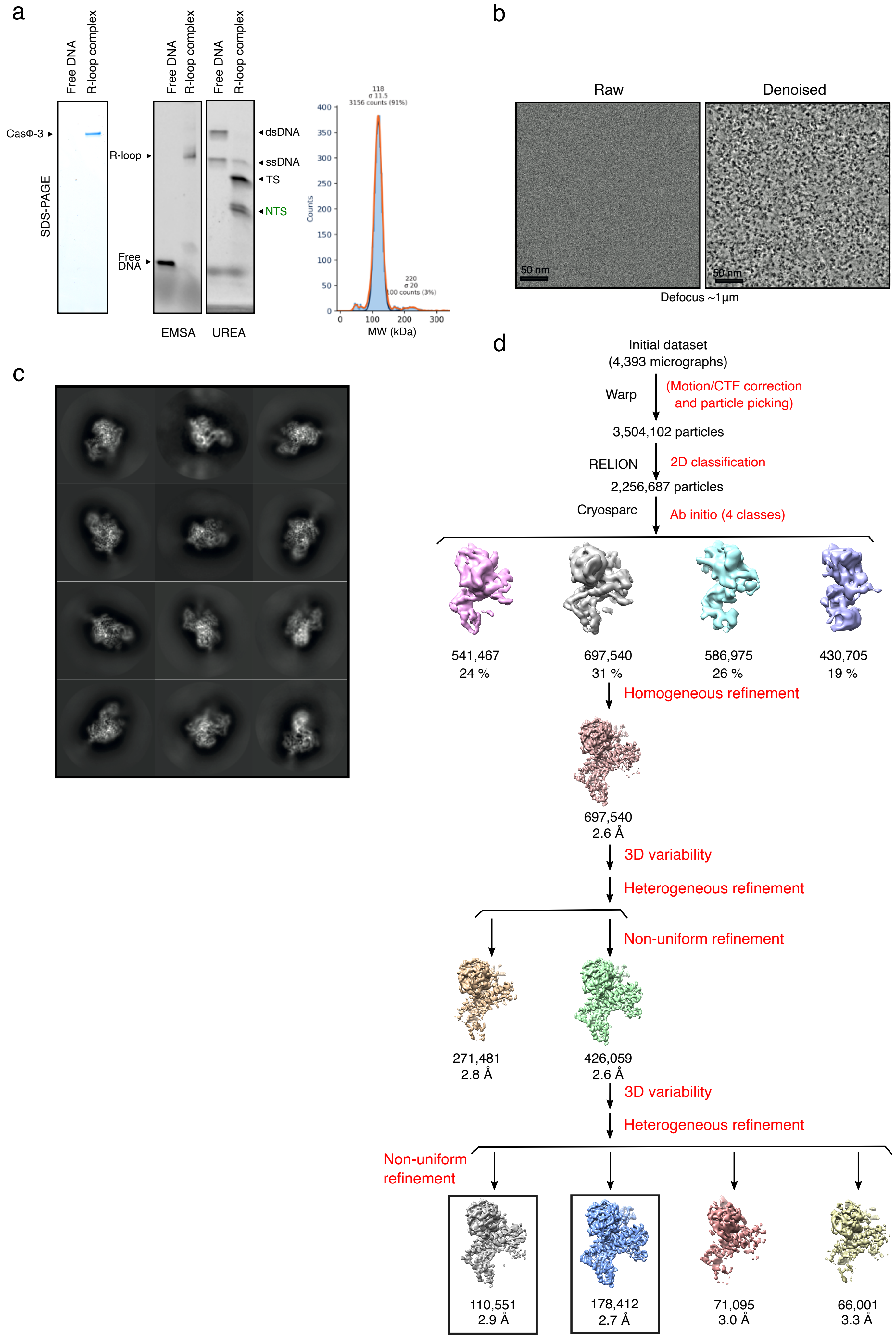
